# Elucidating Molecular Mechanism and Chemical Space of Chalcones through Knowledge Graph and Machine Learning: A Combined Bio-Cheminformatic Pipeline

**DOI:** 10.1101/2024.05.05.592581

**Authors:** Ajay Manaithiya, Ratul Bhowmik, Satarupa Acharjee, Sameer Sharma, Sunil Kumar, Mohd. Imran, Bijo Mathew, Seppo Parkkila, Ashok Aspatwar

## Abstract

We developed a bio-cheminformatics method, exploring disease inhibition mechanisms using machine learning-enhanced quantitative structure-activity relationship (ML-QSAR) models and knowledge-driven neural networks. ML-QSAR models were developed using molecular fingerprint descriptors and the Random Forest algorithm to explore the chemical spaces of Chalcones inhibitors against diverse disease properties, including antifungal, anti-inflammatory, anticancer, antimicrobial, and antiviral effects. We generated and validated robust machine learning-based bioactivity prediction models ((https://ashspred.streamlit.app/) for the top genes. These models underwent ROC and applicability domain analysis, followed by molecular docking studies to elucidate the molecular mechanisms of the molecules. Through comprehensive neural network analysis, crucial genes such as AKT1, HSP90A1, SRC, and STAT3 were identified. The PubChem fingerprint-based model revealed key descriptors: PubchemFP521 for AKT1, PubchemFP180 for SRC, PubchemFP633 for HSP90, and PubchemFP145 and PubchemFP338 for STAT3, consistently contributing to bioactivity across targets. Notably, chalcone derivatives demonstrated significant bioactivity against target genes, with compound RA1 displaying a predictive pIC_50_ value of 5.76 against HSP90A and strong binding affinities across other targets. Compounds RA5 to RA7 also exhibited high binding affinity scores comparable to or exceeding existing drugs. These findings emphasize the importance of knowledge-based neural network-based research for developing effective drugs against diverse disease properties. These interactions warrant further *in vitro* and *in vivo* investigations to elucidate their potential in rational drug design. The presented models provide valuable insights for inhibitor design and hold promise for drug development. Future research will prioritize investigating these molecules for *mycobacterium tuberculosis*, enhancing the comprehension of effectiveness in addressing infectious diseases.

## 1. Introduction

Chalcones, a specialized subclass within the flavonoid family of organic compounds, exhibit a unique structural arrangement, characterized by three aromatic rings (A, B, and C) linked through an α, β-unsaturated carbonyl system(Ferrer et al., 2008; K. Sahu et al., 2012). This distinctive arrangement not only defines their chemical identity but also plays a pivotal role in their diverse biological activities. These naturally occurring compounds, found in various plant species, have piqued the interest of the medicinal chemistry community due to their multifaceted biological actions(Mahapatra et al., 2015).

Structurally, chalcones are composed of three aromatic rings, known as rings A, B, and C. The linkage between the B and C rings through a conjugated double bond system, particularly an α, β-unsaturated carbonyl group, is a defining feature of these compounds(Naik et al., 2020). As members of the flavonoid family, chalcones are essentially plant-derived polyphenolic compounds. Research has highlighted their broad spectrum of cytoprotective and regulatory functions, which are deemed crucial in managing various diseases. A key factor in leveraging chalcones for human use is a thorough understanding of their activity and potential toxicity. The structural conformation of chalcones not only imparts them with specific properties but also contributes to their wide range of biological activities. These include potential roles in reducing inflammation and combating cancer, malaria, *mycobacterium tuberculosis*, and microbial infections(Dhaliwal et al., 2022; Lawrence, 2009; Sivakumar et al., 2007; Ventura et al., 2015). The growing interest in their antimycobacterial capabilities further indicates their potential to develop new treatments for infectious diseases.

The pharmacophoric characteristics of chalcones, which are linked to their chemical composition, are crucial in their use for various health-related applications. The α, β-unsaturated carbonyl group of chalcones is a key feature that facilitates their interactions with biomolecules. As a Michael acceptor, it enables nucleophilic additions with cellular targets like enzymes and receptors, influencing various biological processes such as enzyme inhibition and modulation of signal transduction pathways. The lipophilic nature of chalcones, due to the two aromatic rings (A and C), aids in their interaction with hydrophobic pockets of target proteins, affecting their binding affinity and specificity. Furthermore, the presence of conjugated double bonds between the B and C rings contributes to the absorption of visible light, making chalcones colourful compounds. This aspect also affects their electronic properties and stability. The carbonyl group in the α, β-unsaturated system acts as an electron-withdrawing group, enhancing the reactivity of the double bond and facilitating further interactions with cellular nucleophiles(Mphahlele, 2021; Thapa et al., 2021; Zainuri et al., 2023).

Chalcones with specific substitutions on their aromatic rings have shown significant biological effects. For instance, chalcones with a trifluoromethyl group in ring B and a 3,4,5-trimethoxy substitution in ring A have displayed strong antiproliferative effects against various cancer cells(Karthikeyan et al., 2014; Thapa et al., 2023). Chalcone derivatives with specific substitutions have also shown notable antifungal activity against yeast strains. The antibacterial potential of chalcones varies with the hydrophobic nature of the alkyl chain, indicating that compounds with a medium level of hydrophobicity exhibit potent antibacterial activity. Furthermore, chalcones with trimethoxy substituents and a monofluoro substitution on the B ring have demonstrated enhanced inhibitory activity. The study of chalcones has also extended to their anti-inflammatory efficacy. The presence of electron-withdrawing groups (EWGs) in chalcones has been linked to enhanced antiinflammatory activity. The specific position of the substitution on the phenyl ring A also influences this activity. Chalcone derivatives containing halogens, like fluoride or chloride, have shown significant potential in this regard(Burmaoglu et al., 2017; Goss et al., 1975; Kotra et al., 2010). In the context of antihyperglycemic activity, chalcones with specific substitutions have exhibited notable effects. For example, chalcones with chloro, bromo, iodo, and hydroxy substitutions at certain positions on the A-ring have shown high anti-hyperglycemic activity. Alkyl substitutions on the benzene ring have also improved these effects. The potential of chalcones in treating tuberculosis has become a recent focus of interest. Certain chalcones have demonstrated efficacy against *Mycobacterium tuberculosis* and other related species, making them promising candidates for new drug development, especially for severe TB cases (Chiaradia et al., 2012; Mishra & Jana, 2023; Rozmer & Perjési, 2014).

The study employed a three-step approach to explore chalcone-based small molecules and their interactions with biological systems. The first phase involved using knowledge-based neural networks in a polypharmacology analysis. This allowed us to identify crucial biological processes and genes affected by these compounds, offering a broad view of their impact on different biological pathways. After that, we utilized the same knowledge-based neural networks to predict the bioactivity (measured as pIC_50_ values) of the chalcones derivatives against the identified genes. The study utilizes a Random Forest machine learning algorithm to create ML-QSAR models. This sophisticated machine learning tool enhances the precision of predictions by integrating domain-specific knowledge into the algorithmic learning process. In the final stage, molecular docking studies were conducted to elucidate the structure-function relationships and molecular mechanisms of the chalcones. This provided valuable insights into how these molecules interact with specific genes and biological processes, uncovering potential mechanisms of action. In future efforts, the aim will be to expand the research focus on studying the efficacy of these molecules specifically for tuberculosis treatment. This will involve further exploration of their interactions with *Mycobacterium tuberculosis* and understanding their potential mechanisms of action in combating tuberculosis infections. Additionally, we plan to conduct safety studies of chalcones using the zebrafish larval model and perform *in vitro* and *in vivo* studies using *M. marinum* and zebrafish to validate the safety and effectiveness of chalcone derivatives as potential anti-tuberculosis agents (Aspatwar et al., 2017, 2018; Aspatwar, Hammaren, et al., 2019; Aspatwar, Kairys, et al., 2019).

## 2. Methods and materials

### 2.1. Data Set Collection

Figure 1 illustrates the chemical structures of the chalcone-based compounds synthesized in the laboratory(Acharjee et al., 2018; Sengupta et al., 2017).

**Fig. 1.**
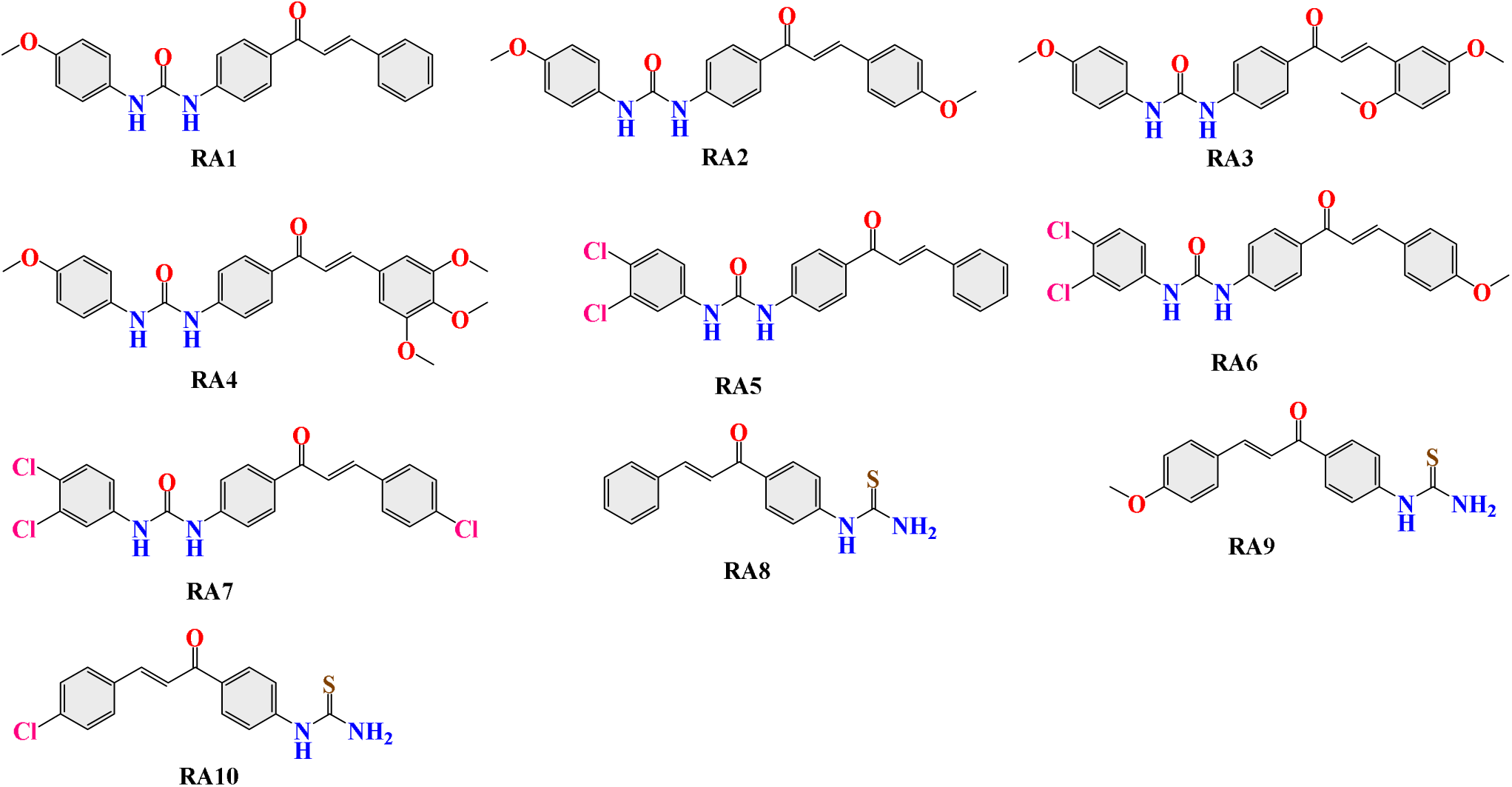
Structural Diagrams of Chalcone-based Derivatives (**RA1-RA10**).

### 2.2. ADME Prediction and Drug Likeliness

To understand the pharmacodynamics of the compounds, it is essential to have a grasp of their physico-chemical properties and pharmacokinetic profile, which includes Absorption, Distribution, Metabolism, and Excretion (ADME). We utilized the SMILES data of the compounds and inputted it into the SwissADME server (https://www.swissadme.ch) to evaluate various physicochemical properties. These properties encompass lipophilicity (iLOGP, XLOGP3, WLOGP, MLOGP, SILICOS-IT, Log P0/w, molar refractivity, topological polar surface area, number of hydrogen bond donors/acceptors), water solubility (Log S - ESOL, Ali, SILICOS-IT), drug-likeness rules (Lipinski, Ghose, Veber, Egan, Muegge), and Medicinal Chemistry (PAINS, Brenk, Lead-likeness, Synthetic accessibility) of the selected compounds(Daina et al., 2017). Moreover, we also acquired important pharmacokinetic parameters for the compounds, such as gastrointestinal (GI) absorption, blood-brain barrier (BBB) permeation, P-gp substrate status, cytochrome-P enzymes inhibition, and skin permeation (logKp), as reported(Pathak et al., 2017). These data provide valuable insights into the behavior and potential of these compounds as pharmacological agents.

### 2.3. Identification of protein targets

The molecular targets of the compounds were determined using the Swiss Target Prediction server (https://www.swisstargetprediction.ch). This server assesses macromolecular targets by comparing 2D and 3D similarities in the active substance library(Pathak et al., 2017). To identify targets associated with diseases, relevant information was retrieved from the Human Gene Database (GeneCards, http://www.genecards.org)(Stelzer et al., 2016) and the Online Mendelian Inheritance in Man (OMIM, http://www.ncbi.nlm.nih.gov/omim) database(Amberger & Hamosh, 2017). The Ven Plot Diagram method was also employed to identify common genes between the compounds and breast cancer. These combined approaches provided valuable insights into potential molecular targets of the compounds and their relevance to different target genes.

### 2.4. Construction of protein-protein interaction

The research involved importing the shared targets of compounds and the disease into the STRING database (https://string-db.org/)(Szklarczyk et al., 2021) to construct protein-protein interactions (PPI) within Homo sapien. Solid circles represent genes in the resulting PPI network, and the enclosed structures represent the corresponding proteins. The genes are interconnected by lines of various colors, indicating the biological processes between the proteins. The researchers utilized the Cytoscape plug-in (v3.8.0) and CytoHubba (http://www.cytoscape.org/) to visualize the interaction network among targets and compounds. In this network, nodes represent compounds and targets, while edges represent interactions between compounds and target(Shannon et al., 2003).

### 2.5. Gene Ontology (GO) Analysis

FunRich 3.1.3 was utilized for functional enrichment analysis of gene ontology, encompassing biological pathways (BPA), cellular components (CC), and biological processes (BPR)(Fonseka et al., 2021). Moreover, the researchers employed the Kyoto Encyclopedia of Genes and Genomes (KEGG) pathways to identify pathways associated with breast cancer among the selected targets. A p-value lower than 0.05 was considered statistically significant and indicative of the relevance of the gene ontology analysis.

### 2.6. Machine Learning Assisted QSAR Study

#### 2.6.1. Data Collection and Pre-Processing

The **panda’s** library efficiently manipulates structured data, including data frames. The **chembl_webresource_client** library is designed to access the extensive ChEMBL database, particularly for bioactive molecules and biological activities related to genes AKT1, SRC, HSP90A, and STAT3. From the ChEMBL database, a collection of inhibitors was obtained: 4170 for AKT1, 5172 for SRC, 1369 for HSP90A, and 1437 for STAT3, along with their corresponding IC_50_ values. Biological activity (IC_50_) of molecules is categorized as active (below 1000 nM), intermediate (1000-10000 nM), or inactive (above 10000 nM). Exploratory data analysis or chemical space analysis was conducted to traverse the chemical landscape of inhibitory compounds using Lipinski’s rule of five descriptors: Molecular weight, ALogP, hydrogen bond donor, and hydrogen bond acceptor.

##### Machine Learning-Based QSAR Modeling Process

All modeling processes are done using the python programming language in Google Colab, facilitated by the Scikitlearn package (version 1.0.2).

#### 2.6.2. Molecular Fingerprints Calculation

PubChem fingerprints provided by the PaDEL package (PaDELpy-0.1.13) were used for modelling(Yap, 2011). The fingerprint set contains 881 binary representations of the chemical structural fragments used by PubChem. The parameter for the PaDEL package is set to detect aromaticity: true; standardize nitrogen: true; standardize tautomers: true; threads = 2; remove salt: true; log = true; fingerprints = true.

#### 2.6.3. Feature Selection

Features with variance lower than 0.1 and features demonstrating high correlation (>0.95) were removed. As a result of AKT1, after feature selection of the 881 features, there are 213 lefts after removing low-variance features and high correlation features. As a result of SRC, after feature selection of the 881 features, there are 241 lefts after removing low-variance features and high-correlation features. As a result of HSP90A, after feature selection of the 881 features, there are 253 lefts after the removal of low-variance features and high correlation features. As a result of STAT3, after the feature selection of the 881 features, there are 262 lefts after removing low-variance features and high-correlation features.

#### 2.6.4. QSAR Model Construction

For all four models, the ratio of the training set and testing set is set to 80:20. Using the Kennington Stone algorithm, the final best AKT1, SRC, HSP90A, and STAT3 inhibitor from the ChEMBL dataset was used to divide the dataset into training and test sets with an 80:20 split ratio (*DTClab.Https://Dtclab.Webs.Com/Software-Tools*; *Github.Https://Github.Com/Dataprofessor/Code/Tree/Master/Python*;*Padel.Http://Www.Yapcwsoft.Com/Dd/Padelescriptor/*). A user-defined variance cut-off value was employed to retrieve significant descriptors from the ChEMBL dataset, and constant descriptors with identical or nearly identical values for all compounds were eliminated. Similarly, inter-correlated descriptors were identified and processed according to the intercorrelation coefficient cut-off value specified by the user.

#### 2.6.5. Development and Validation of Random Forest based QSAR models

The datasets from ChEMBL (AKT1, SRC, HSP90A, and STAT3) were employed in the creation of Random Forest (RF)-based machine learning models using Weka Software(*Weka.Https://Www.Cs.Waikato.Ac.Nz/Ml/Weka/*). RF, a supervised machine learning technique, encompasses a collective ensemble of predictors that is fundamentally derived from decision trees. Training of RF is carried out using the Bagging or Bootstrap aggregation technique. Additionally, the RF algorithm attempts to tackle the issue of overfitting commonly associated with decision trees. Subsequently, a correlation coefficient (R2) comparison between the training and test datasets was utilized to validate the Quantitative Structure-Activity Relationship (QSAR) models. The most valuable features were identified through the application of the RF Regressor algorithms for the RF models, which were then depicted in Variance Importance Plots (VIP). A graphical comparison of experimental versus predicted values for each QSAR model was conducted using the matplotlib Python package(*Github. Https://Github.Com/Vappiah/Machine-Learning-Tutorials*).

Furthermore, the construction of receiver operating characteristics (ROC) graphs for both QSAR models was carried out with a pre-existing Python script designed for multi-class model classification. The Receiver Operating Characteristic (ROC) acts as a graphical instrument for assessing the performance of classifiers and evaluating the effectiveness of classification-based Quantitative Structure-Activity Relationship (QSAR) models by leveraging the features derived from the confusion matrix. It provides a two-dimensional depiction (0 to 1) of the true positive (TP) rate contrasted against the false positive (FP) rate(Pedregosa et al., 2012). The area under the curve (AUC) was formulated as a quantitative metric to provide a competitive evaluation of ROC analysis. The AUC of a ROC plot stands as a reliable proxy for a discriminant model’s performance, with its value ranging from zero (total misclassification) to one (perfect classification).

Moreover, the applicability domain (AD) of both QSAR models was assessed through the bounding box technique of principal component analysis (PCA). This requires a PCA examination of the scores plot to compare the molecules’ chemical space from the training and test sets(Sahigara et al., 2012). The AD was ascertained using the PCA function from the sklearn—decomposition module of the scikit-learn machine learning toolkit in Python(*Scikit-Learn. Https://Github.Com/Scikit-Learn/Scikit-Learn.Git.*).

### 2.7. Molecular Docking

A molecular docking approach was employed to evaluate the inhibitory potential of chalcone derivatives across various activities, including antibacterial, anticancer, antidiabetes, anti-inflammation, and antifungal effects. The protein structures pertinent to the investigation, namely AKT1 (PDB ID: 4EJN)(Ashwell et al., 2012), SRC (PDB ID: 2OIQ)(Seeliger et al., 2007), HSP90A (PDB ID: 3O0I)(Patel et al., 2013), and STAT3 (PDB ID: 6NJS)(Bai et al., 2019), were sourced from the Protein Data Bank in PDB format. Active sites within each protein structure were pre-delineated to facilitate docking by constructing grid boxes around the co-crystallized ligand. The AutoDock Tools software(Trott & Olson, 2010) was then employed to prepare the protein molecules. This process involved rectifying missing residues, eliminating water molecules, adding polar hydrogens, and applying Kollman charges. The resulting protein structures were saved in pqbqt format. Ligand molecules’ 2D structures underwent conversion to 3D structures using the MMFF94 force field within the AutoDock Vina software. These transformed ligand structures were saved and converted to pdbqt format utilizing the Open Babel GUI. In the final step, a Perl script, in conjunction with Perl software, facilitated the docking of all ligand molecules against the protein structures. The resulting binding affinities or docking scores for each ligand molecule and respective target receptor were quantified in kcal/mol units. To glean insights into the molecular interactions, Pymol and Discovery Studio Visualizer were employed, enabling an in-depth exploration of ligand binding interactions with the most favorably binding proteins.

### 2.8. Molecular Dynamic

To explore the stability of the most promising molecule in biological conditions, we carried out molecular dynamics (MD) simulations. These simulations are essential for understanding how the molecule behaves in a solvent environment. We set up the simulation in an orthorhombic box with dimensions of 12 Å on each side, using the buffer size method to optimize the volume of the box. The simulations were conducted using the TIP3P water model and the OPLS3e force field by Schrodinger Inc., which are standards for simulating proteins and ions. Sodium chloride was added to the system at a concentration of 0.15M to mimic physiological conditions, with sodium (Na+) and chloride (Cl-) ions. The simulations ran for 100 nanoseconds using the Desmond Molecular Dynamics module, producing around 1000 snapshots of the system’s behavior. These were performed under the NPT ensemble, maintaining a constant temperature of 300 K and a pressure of 1 bar, ensuring the system was equilibrated before the simulations began.

## 3. Results

### 3.1. ADME Prediction and Drug Likeliness

ADME detection is crucial in drug discovery and development. Analyzing structural and physicochemical characteristics helps identify compounds with favorable pharmacokinetic profiles and drug-like features, minimizing drug-drug interactions and experimental failures. SwissADME databases provide efficient models to predict compound properties, aiding in drug development decision-making.

The physicochemical properties, pharmacokinetic profile, and medicinal characteristics of the selected compounds are depicted in **Table 1**. **See Supplementary file S1** for more information on **Table 1**. All combinations shared an identical bioavailability score of 0.55, indicating moderate bioavailability. None of the compounds exhibited any PAINS alerts, suggesting a lack of common structural motifs associated with assay interference. Most compounds showed inhibitory activity against various cytochrome P450 (CYP) enzymes, potentially affecting drug metabolism and interactions. However, all compounds were predicted as nonsubstrates of P-glycoprotein (Pgp), reducing the risk of drug-drug interactions mediated by this efflux transporter. All compounds demonstrated high gastrointestinal (GI) absorption, indicating efficient absorption in the gastrointestinal tract. However, none of the compounds were predicted to permeate the blood-brain barrier (BBB).

**Table 1:**
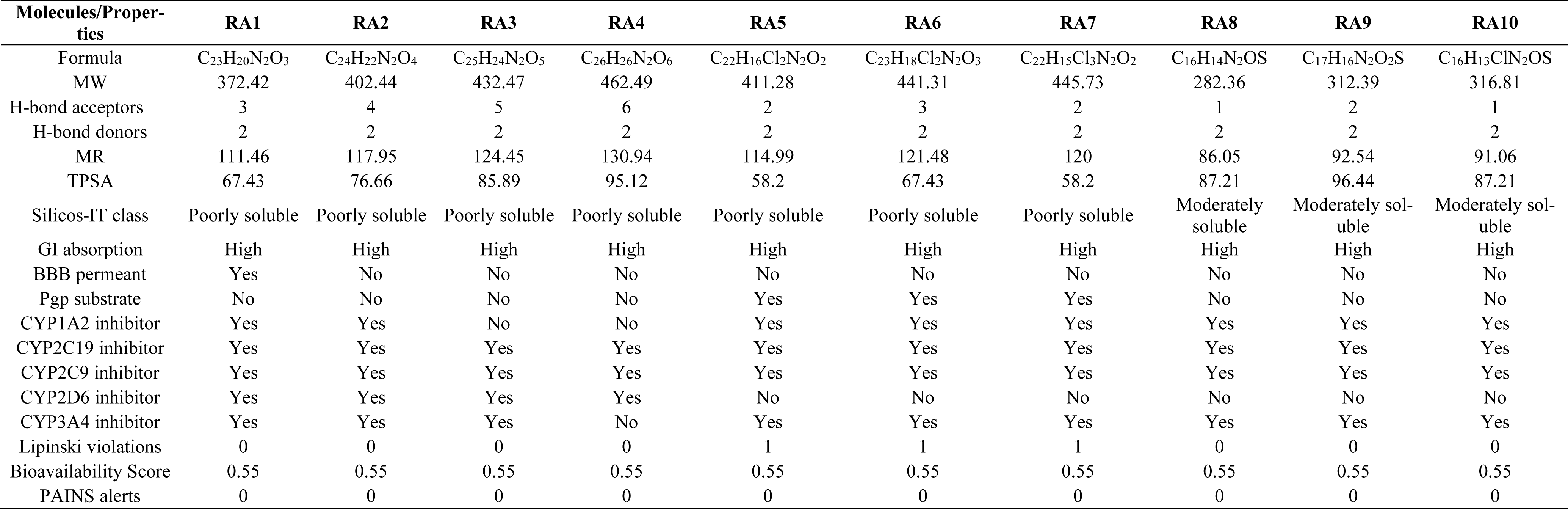
ADME parameters of the compounds RA1-RA10.

All compounds adhered to Lipinski’s Rule of Five, suggesting favorable drug-like properties concerning absorption and distribution. However, based on the Silicos-IT class, most compounds were classified as poorly soluble, with Compound RA8 being moderately soluble. **Compound RA4** displayed the highest molar refractivity (130.94), suggesting its potential for intermolecular interactions and polarizability. On the other hand, **Compound RA8** had the lowest molar refractivity (86.05). Regarding the topological polar surface area, **Compound RA4** exhibited the highest value (95.12 Å²), while **Compound RA5** displayed the lowest value (58.2 Å²). These properties influence a compound’s solubility and permeability characteristics.

### 3.2. Compound and disease targets

We obtained compound targets from the Swiss target prediction server and marks associated with four diseases from the Human gene and OMIM databases (**See Supplementary file, s1**). The Venn diagram demonstrates the intersection of 346 targets related to bacterial diseases, 346 targets associated with inflammation, 364 targets linked to cancer, 349 targets relevant to diabetes, and 220 targets about fungal diseases. These intersections represent the common gene targets shared between compounds and the specified diseases (**see Fig. 2**.).

**Fig. 2.**
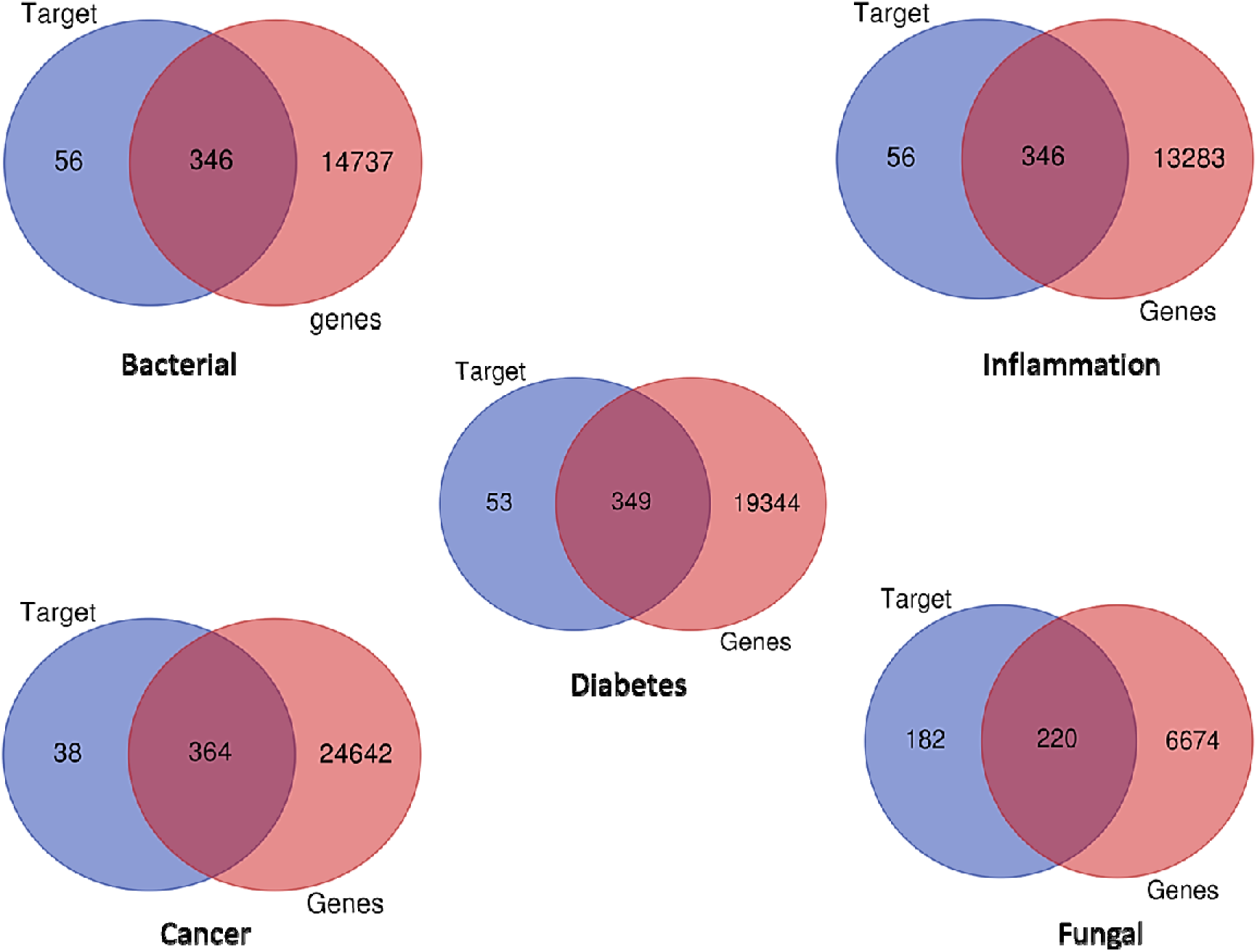
Overlapping targets between the potential compound’s targets and disease-related genes using Ven Plot Diagram.

### 3.3. PPI Network analysis

The PPI network was constructed using the STRING database, explicitly focusing on targets from Homo sapiens. This research compares five biological networks: bacterial, cancer, diabetes, fungal, and inflammation. The networks were constructed using data from the STRING database and visualized in Cytoscape after importing the data in a .tsv format(Lopes et al., 2010). For each network, the number of nodes and edges was recorded: bacterial network (345 nodes, 4342 edges), cancer network (363 nodes, 4555 edges), diabetes network (348 nodes, 4412 edges), fungal network (220 nodes, 2814 edges), and inflammation network (345

Key network properties were calculated and analyzed for each network. These properties included the average local clustering coefficient, PPI enrichment p-value, and average node degree. The bacterial network had an average local clustering coefficient of 0.458, a PPI enrichment p-value of p<1.0e-16, an average node degree of 25.2, and an Average Shortest Path Length of 2.46. Similarly, the cancer network had an average local clustering coefficient of 0.459, a PPI enrichment p-value of p<1.0e-16, an average node degree of 25.1, and an Average Shortest Path Length of 2.48. The diabetes network had an average local clustering coefficient of 0.462, a PPI enrichment p-value of p<1.0e-16, an average node degree of 25.4, and an Average Shortest Path Length of 2.46. The fungal network had an average local clustering coefficient of 0.52, a PPI enrichment p-value of p<1.0e-16, an average node degree of 25.6, and an Average Shortest Path Length of 2.26. The inflammation network had an average local clustering coefficient of 0.457, a PPI enrichment p-value of p<1.0e-16, an average node degree of 25.8, and an Average Shortest Path Length of 2.44.

Furthermore, the networks’ structural characteristics were analyzed, including the network diameter and radius. The bacterial, cancer, and inflammation networks had a diameter of 6 units and a radius of 3 units. Diabetes and cancer networks had a diameter of 6 units and a radius of 4 units.

Fig. 3A and 3B display the compound-target interactions that were constructed using Cytoscape. The top 10 targets were subjected to network analysis, and the degree of freedom for each target was reported in **Table 2** and Fig. 3B. Among the ten genes shared across the five diseases, AKT1, SRC1, HSP90AA, and STAT3 exhibited strong associations in all five diseases. These four genes were selected based on their degree scores, ranging between 91 and 148. In each disease, AKT1, SRC1, HSP90AA, and STAT3 displayed the highest degrees, surpassing 110, except for fungal disease, where their degrees were above 90. Notably, AKT1 demonstrated prominent significance by securing the first rank in bacterial, inflammation, and cancer networks, with 148, 149, and 149 scores, respectively. In the fungal and diabetes networks, AKT1 remained highly significant, achieving the first rank with scores of 119 and 148, respectively. AKT1, a serine/threonine kinase, is intricately involved in diverse cellular processes such as cell survival, proliferation, and metabolism(Schiliro & Firestein, 2021). In bacterial infections, AKT1 signaling has been associated with the modulation of host immune responses and pathogen invasion mechanisms. In cancer, AKT1 dysregulation is frequently seen, leading to tumor growth and reduced responsiveness to conventional treatment methods. Additionally, AKT1’s critical role in glucose metabolism and insulin signaling makes it an attractive target in diabetes management. AKT1 signaling modulates antifungal immune responses, and its inhibition has shown potential in enhancing the host’s ability to combat fungal infections(Fayard et al., 2010).

**Figure 3A.**
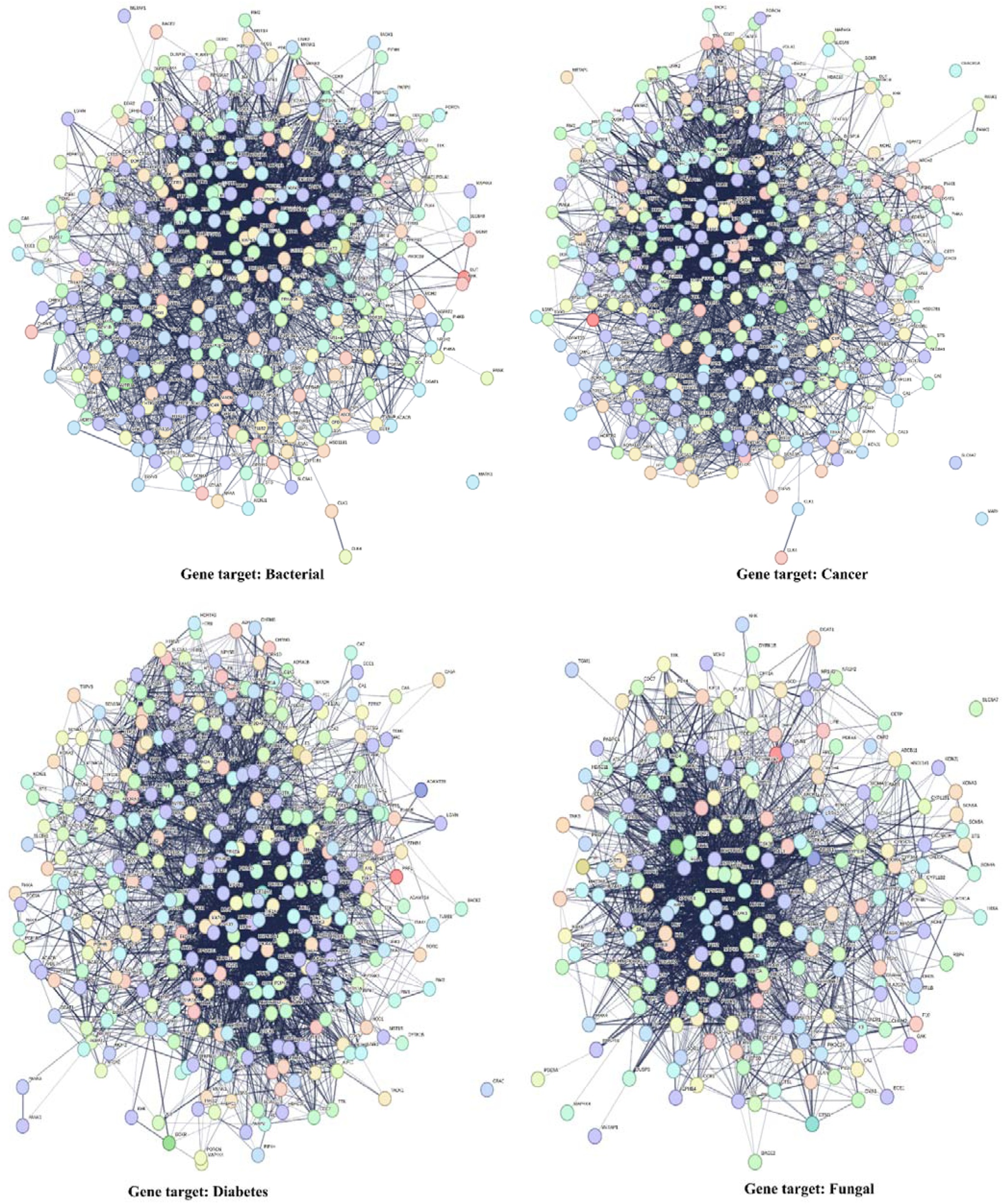
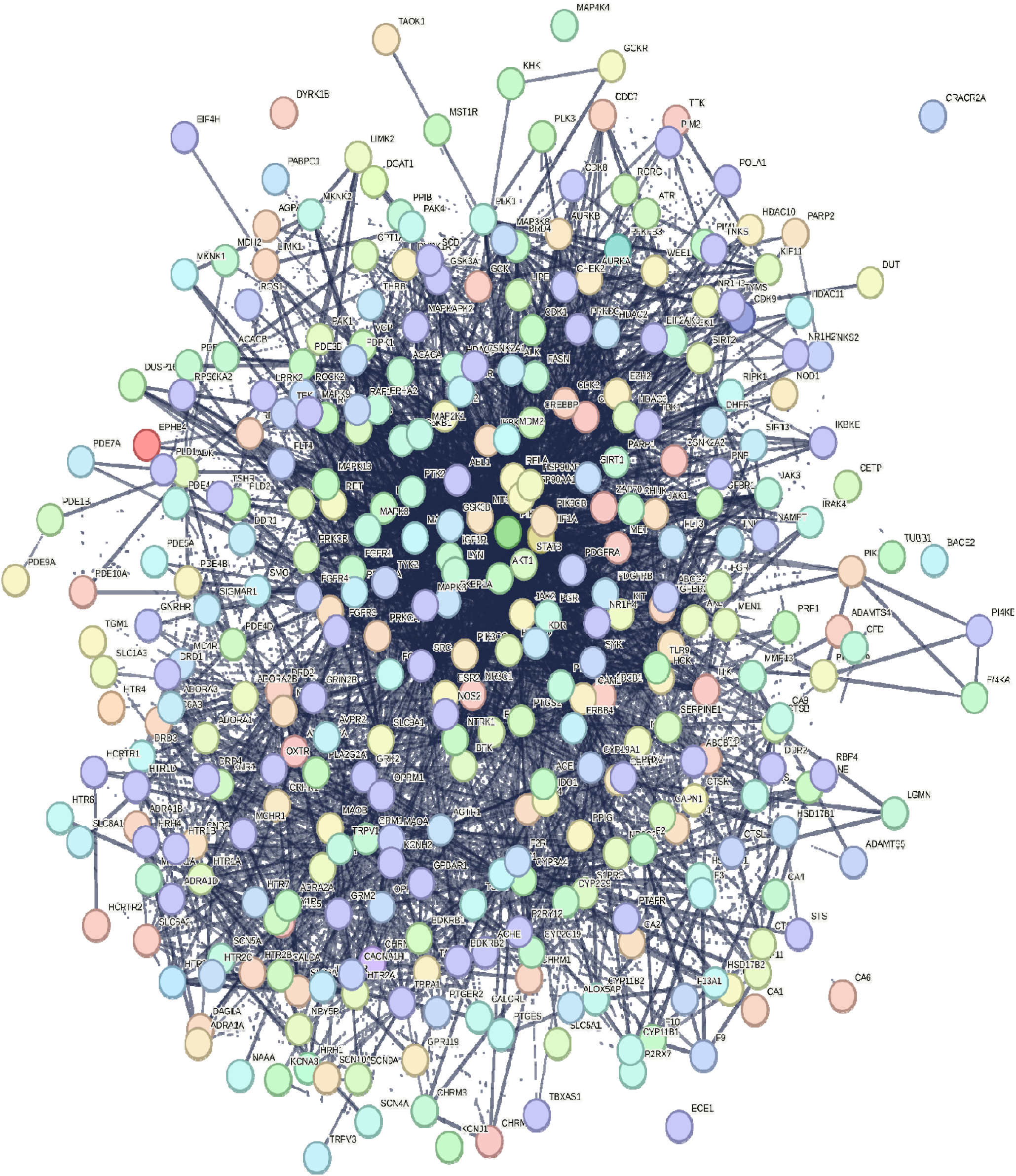
Interactions of gene targets in four diseases (Bacterial, Cancer, Diabetes, and Fungal) were visualized using Cytoscape and Network Analysis.

**Figure 3B.**
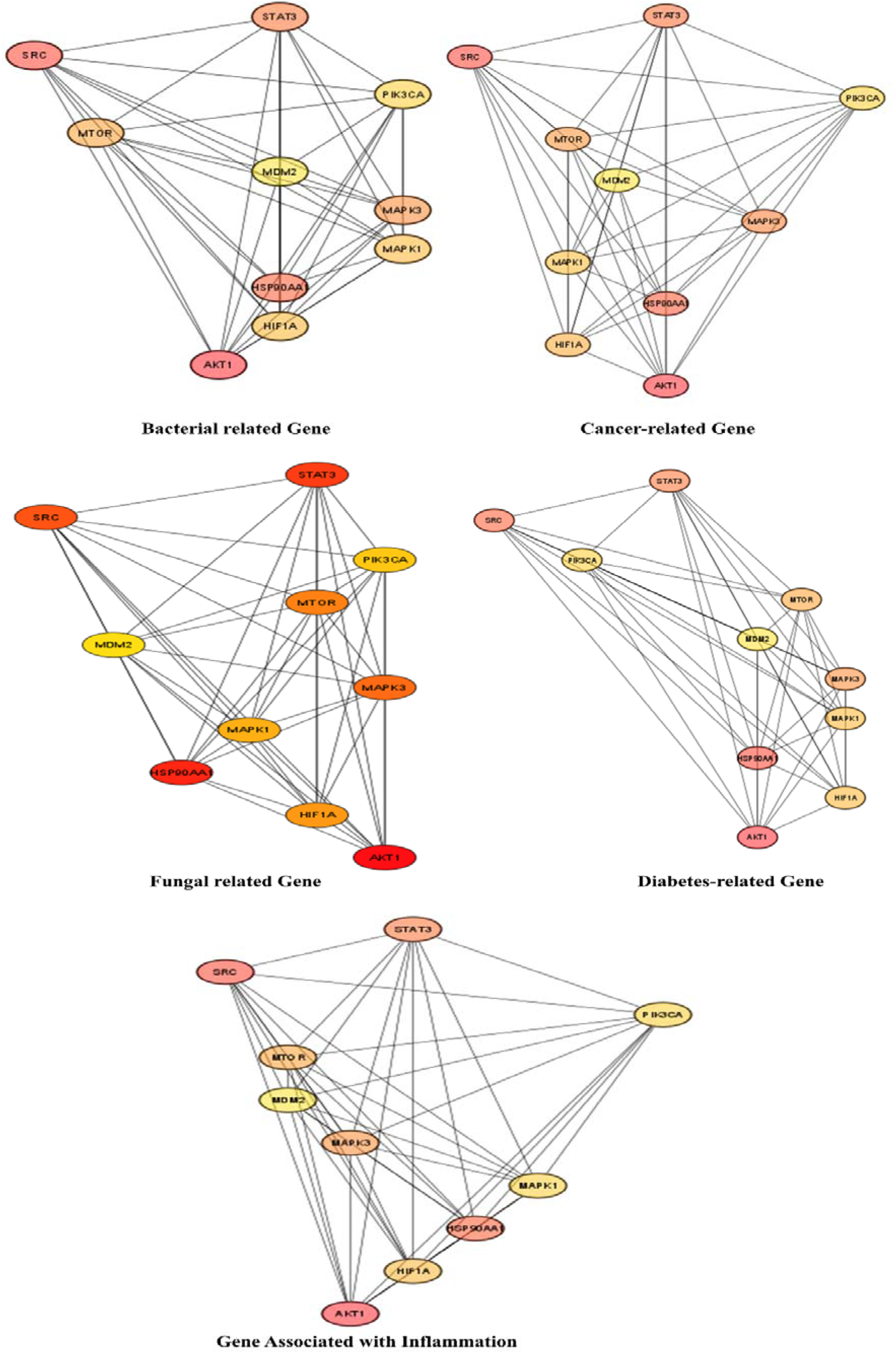
Top 10 Gene Target Interactions in Four Diseases Visualized through Cytoscape and Analyzed Using Network Analysis.

**Table 2.**
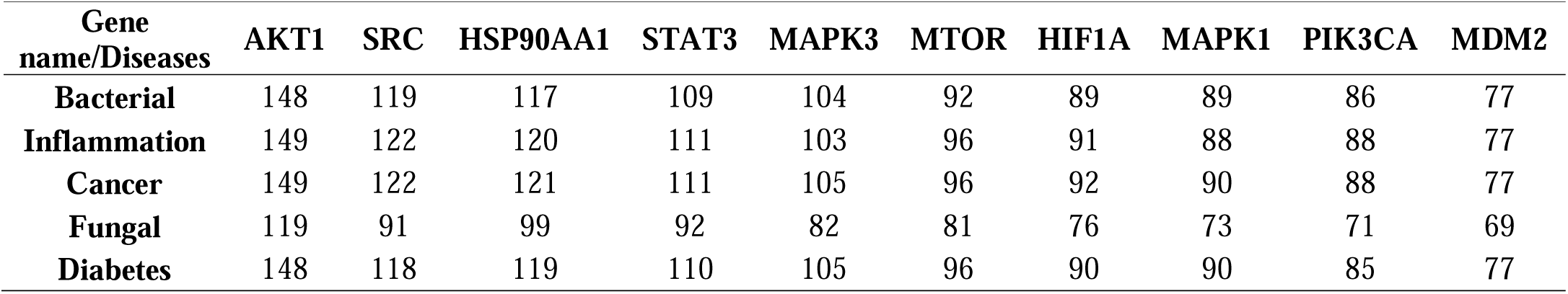
Degree of freedom of top 10 gene and their scores.

SRC, a non-receptor tyrosine kinase, plays a pivotal role in signal transduction pathways that govern cell growth, motility, and invasion. In bacterial infections, SRC has been linked to host cell invasion and the intracellular survival of pathogen(Siddiqui et al., 2012). In the realm of cancer, SRC is frequently overexpressed, promoting tumor progression and metastasis. Furthermore, SRC participates in insulin signaling and glucose metabolism, making it relevant to diabetes research. Additionally, SRC activation is involved in inflammation-related processes, contributing to the pathogenesis of various diseases(S. T. Liu et al., 2014).

SRC, a key cellular motility and adhesion regulator, contributes to fungal invasion and dissemination. Targeting SRC using specific inhibitors may impede fungal spread and improve treatment outcomes. HSP90AA1, a heat shock protein, is a molecular chaperone that plays a pivotal role in protein folding and stability. In bacterial infections, HSP90AA1 facilitates bacterial virulence by promoting the stability of bacterial effectors.

Within the cancer domain, HSP90AA1 acts as a critical chaperone for oncoproteins and proteins associated with drug resistance(Backe et al., 2020). Its implication in insulin resistance and β-cell dysfunction highlights its relevance in diabetes research. Moreover, HSP90AA1 is involved in inflammatory responses across various diseases. HSP90AA1 functions as a chaperone for fungal proteins, essential for fungal survival and virulence. Disrupting HSP90AA1’s function has been explored as a strategy to weaken fungal pathogens. STAT3, a transcription factor, is essential in cell survival, proliferation, and immune responses(Yu et al., 2009). STAT3 signaling modulates the host’s inflammatory and immune responses to bacterial invasion in bacterial infections. STAT3 activation promotes tumor growth and immune evasion in the context of cancer. In diabetes research, STAT3 influences insulin signaling and pancreatic β-cell function. STAT3 mediates inflammation-related processes, impacting disease pathogenesis. STAT3 plays a vital role in orchestrating immune responses during fungal infections, and inhibiting its activity might enhance the host’s antifungal defense mechanisms(Y. C. Liu et al., 2019; Vella et al., 2023). The analysis of protein ranking across diverse biological networks offers valuable insights into the relative significance of AKT1, SRC, HSP90AA1, and STAT3 in various cellular processes and disease contexts. The consistently high rankings of these proteins suggest their crucial roles in cellular regulation, signal transduction, and disease development. Based on the Cytohubba analysis, all synthesized compounds exhibited the highest score, indicating their interactions with the maximum number of identified elements in all five diseases, achieving a score of 100. Previous research has highlighted the potential of chalcone-based novel phenyl ureas as effective antihyperglycemic agents with a likely PPAR gamma agonistic action.

### 3.4. Gene Ontology

We performed a functional enrichment analysis using the FunRich software on the top 10 targets selected based on their degree. However, the degree of gene targets has different ranks; all targets, diabetes, inflammation, fungal, bacterial, and cancer, have almost the same 10 ten degrees of genes identified. Based on the data analysis of target genes, all diseases have the same cellular component, biological pathway, and process. Fig. 4 illustrates the top 10 Biological Pathway Annotations, Cellular Component Annotations, and Biological Process Annotations. Among the top 10 biological pathways identified, the following pathways were found: NGF signaling via TRKA from the plasma membrane 80%, Signaling by EGFR 70%, Signaling by FGFR 70%, ErbB2/ErbB3 signaling events 60%, Signaling by PDGF 70%, Downstream signal transduction 70%, Signalling by NGF 80%, Signaling by SCF-KIT 70%, VEGFR1 specific signals 60%, IL2-mediated signaling events 80%.

**Fig. 4.**
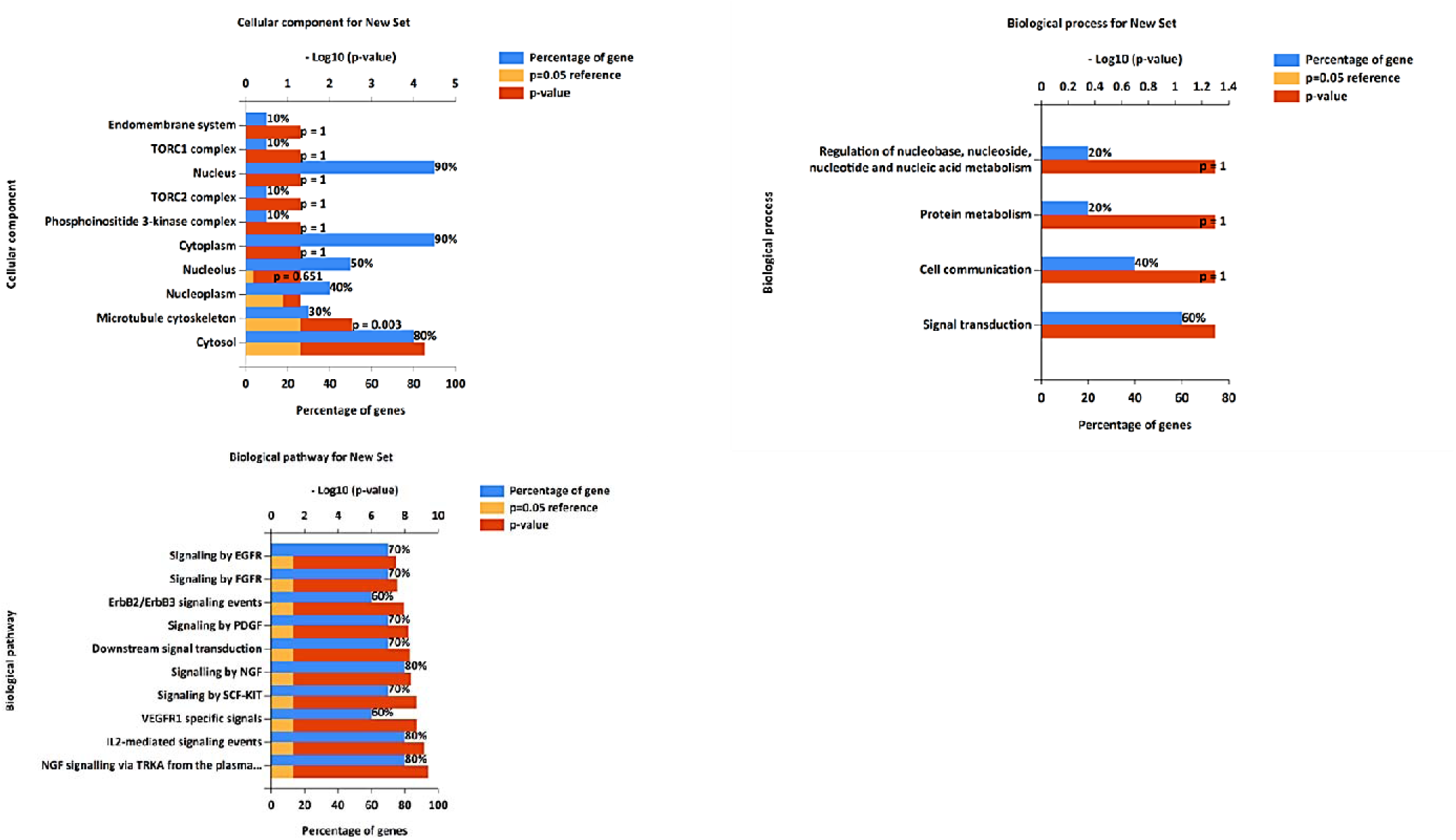
Gene ontology analysis: Cellular components, biological processes, and biological pathway.

The scientific literature has extensively discussed that these pathways are involved in diabetes, inflammation, fungal, bacterial, and cancer development. Nerve Growth Factor (NGF) is a neurotrophic factor involved in neurons’ development, survival, and function. Enriching genes in this pathway suggests their potential roles in mediating NGF signaling through its receptor TRKA (NTRK1). SRC, STAT3, and MAPK1, in particular, are known to be involved in neuronal signaling and synaptic plasticity, and they may play important roles in the downstream events of NGF-TRKA signaling(Chao, 2003). Interleukin-2 (IL-2) is a cytokine central to regulating immune responses. Enriching genes in this pathway suggests their potential roles in mediating IL-2 signaling events. SRC and STAT3 are known to be involved in immune cell signaling and activation.

In cancer, IL-2 has been used as an immunotherapy to stimulate the immune system’s anti-tumor response, and SRC and STAT3 may be involved in the downstream effects of IL-2-mediated immune activation(Rosenberg & Restifo, 2015). EGFR signaling is closely linked to various types of cancers, including lung cancer, breast cancer, colorectal cancer, and head and neck cancer. Dysregulation of EGFR, such as overexpression or activating mutations, can lead to uncontrolled cell proliferation, invasion, and metastasis in these malignancies(Lemmon & Schlessinger, 2010). EGFR signaling, while not a central factor in the development of diabetes, may impact certain cellular responses associated with complications of the disease, such as diabetic retinopathy. Similarly, while EGFR signaling doesn’t directly correlate with bacterial or fungal infections, it might indirectly affect the immune responses to these infections. This is possible due to the expression of EGFR in diverse immune cells and tissues, suggesting its involvement in modulating host responses to various health challenges. FGFR (Fibroblast Growth Factor Receptor) is another family of receptor tyrosine kinases involved in cell proliferation, migration, and differentiation. Fibroblast growth factors (FGFs) binding to FGFR leads to receptor dimerization and activation of downstream signaling pathways.

FGFR signaling is critical in development, tissue repair, and angiogenesis. Aberrant FGFR signaling has been implicated in various cancers and developmental disorder(Turner & Grose, 2010). FGFR signaling, while not directly involved in bacterial or fungal infections nor a primary contributor to diabetes development, may have a role in inflammation. However, its explicit involvement in inflammation-associated diseases warrants further study. VEGFR1 signaling plays a crucial role in angiogenesis, forming new blood vessels. It is expressed in tumor cells and various immune cells, making it relevant to several diseases, including cancer, inflammation, diabetes, and vascular diseases. In cancer, particularly colorectal and breast cancer, VEGFR1 signaling contributes to tumor angiogenesis and growth. High levels of VEGFR1 expression are associated with poor prognosis in these cancers. As a result, targeting VEGFR1-specific signals is being investigated as a potential strategy for cancer treatment. In inflammatory diseases like rheumatoid arthritis and inflammatory bowel disease, VEGFR1-mediated signals play a role in recruiting immune cells and promoting angiogenesis to facilitate tissue repair. Consequently, interventions focused on regulating VEGFR1 are under investigation to manage the progression of diabetic retinopathy. VEGFR1 signaling also affects the progression of various vascular diseases, such as atherosclerosis and vascular malformations. It can modulate angiogenesis within atherosclerotic plaques and contribute to abnormal vessel development in vascular malformations(Bollenbecker et al., 2023; Shibuya, 2011).

PDGF (Platelet-Derived Growth Factor) is a growth factor in cell proliferation and wound healing. It signals through two receptor tyrosine kinases, PDGFRα and PDGFRβ. Upon ligand binding, PDGF receptors undergo autophosphorylation and activate downstream signaling pathways, including the PI3K-AKT and MAPK pathways. PDGF signaling is important in tissue repair, angiogenesis, and development. Aberrant PDGF signaling has been implicated in cancer and fibrotic diseases(Heldin & Lennartsson, 2013). While not directly associated with fungal infections, PDGF signaling may be involved in regulating inflammation and tissue repair. Additionally, it may hold relevance for diabetic complications, including nephropathy and retinopathy. The downstream Signal Transduction pathway involves the transmission of signals from activated cell surface receptors (such as EGFR, FGFR, and PDGFR) to intracellular effectors. Downstream signal transduction pathways include MAPK/ERK, PI3K-AKT, and JAK-STAT. These pathways regulate gene expression and modulate cellular responses, such as proliferation, survival, and differentiation. Dysregulation of downstream signal transduction can lead to various diseases, including cancer and inflammatory disorders(Shen et al., 2020). ErbB2 (HER2) and ErbB3 (HER3) are members of the EGFR family of receptor tyrosine kinases. They form heterodimers and activate downstream signaling pathways upon ligand binding or through other mechanisms. ErbB2 does not bind a specific ligand but can enhance signaling by forming heterodimers with other ErbB family members. ErbB2/ErbB3 signaling plays crucial roles in the cell proliferation, survival, and metastasis of various cancers. Abnormal ErbB2 (HER2) expression is closely linked with aggressive forms of breast cancer, and targeted treatments focusing on ErbB2 have demonstrated clinical effectiveness (Hynes & Lane, 2005). SCF (Stem Cell Factor) and KIT (KIT proto-oncogene) are involved in hematopoiesis, melanogenesis, and cell survival. The binding of SCF to its receptor KIT activates down-stream signaling events. KIT signaling is crucial for stem cell development and hematopoiesis—aberrant KIT signaling(Lennartsson & Rönnstrand, 2012).

The proteins SRC, HSP90AA1, STAT3, MAPK3, MTOR, HIF1A, MAPK1, PIK3CA, and MDM2 are involved in several important biological pathways that have relevance to various diseases, including cancer, diabetes, inflammation, and other disorders. These proteins are crucial in signal transduction, growth regulation, immune responses, and cellular metabolism. When these pathways and proteins become dysregulated, they can contribute to the development and progression of diseases. In cancer, these proteins often promote cell growth, survival, and metastasis. Dysregulation of these pathways can lead to uncontrolled cell proliferation and tumor formation. For example, the MAPK pathway (involving proteins like MAPK1 and MAPK3) is frequently altered in cancer, leading to excessive cell division and tumor growth. In diabetes, proteins like MTOR and PIK3CA are involved in insulin signaling and glucose metabolism. Dysfunctional signaling in these pathways can affect insulin sensitivity and glucose regulation, contributing to diabetes and its complications. In inflammation, proteins like STAT3 and HIF1A are key players in immune responses and inflammation regulation. Aberrant activation of these proteins can lead to chronic inflammation associated with various inflammatory diseases.

The gene ontology analysis uncovers critical biological processes associated with the top five targets for diseases, including diabetes, inflammation, fungal, bacterial, and cancer. Notable processes include signal transduction (60%), protein metabolism (20%), cell communication (40%), energy pathways (10%), and regulation of nucleobase, nucleoside, nucleotide, and nucleic acid metabolism (20%) **(See Fig. 4)**. These insights are invaluable, shedding light on the molecular mechanisms driving disease development and progression. Additionally, identified cellular components associated with the top ten disease targets reveal where these elements predominantly exist within cells. These locations include the phosphoinositide 3-kinase complex (10%), TORC1 and TORC2 (10%), the nucleus (90%), nucleoplasm (40%), endomembrane system (10%), the nucleolus (50%), the TORC2 complex (10%), the cytoplasm (90%), microtubules, and the cytosol (80%) (**See** Fig. 4). Understanding the cellular location of these targets provides crucial insights into their functional roles in specific diseases, allowing for more targeted and precise intervention strategies.

### 3.5. KEGG pathway

In the present research, the examined compounds showcased distinctive impacts on conditions such as cancer, diabetes, and inflammation. These substances showed promising effects on ovarian neoplastic cells through their interaction with the MAPK signaling pathways, particularly focusing on the ERK element and the MAPK receptors(Dhillon et al., 2007). This observation is harmonious with the well-established function of MAPK receptors in fostering tumor proliferation and survival. The compounds also impacted the mTOR signaling pathway, targeting elements such as PK13, AKT1, and mTOR receptors, which are crucial for cell proliferation and survival(Saxton & Sabatini, 2017). Similarly, these compounds affected the JAK-STAT signaling pathway, explicitly targeting components like STAT3 receptors(O’Shea et al., 2015). The compounds influenced the MAPK signaling pathway by activating components via the EGFR receptors. Regarding the HIF1 alpha signaling pathway, the compounds’ role is noteworthy. RTKs activate HIF1 alpha, so by targeting them, the compounds could potentially deregulate their activity.

The PI3K-AKT signaling pathway is a crucial intracellular signaling pathway implicated in multiple cellular functions, such as cell growth, proliferation, angiogenesis, and survival. It is activated by various types of cellular stimuli or toxic insults(Manning & Toker, 2017; Porta et al., 2014). The PI3K-AKT pathway is central to insulin signaling. When insulin binds to its receptor, it triggers the activation of PI3K, leading t the activation of AKT. AKT subsequently stimulates glucose uptake by promoting the translocation of the glucose transporter GLUT4 to the cell membrane. This pathway’s alteration can lead to insulin resistance, a key factor in developing type 2 diabetes. Dysregulation in PI3K/AKT signaling has been associated with diabetic complications, including nephropathy and retinopathy(Oeckinghaus & Ghosh, 2009; Sadikot et al., 2005) **(See Fig. 5A)**. In inflammation disease, the activation of the NF-κB pathway, including the resultant upregulation of BCL-XL and c-Myb, can contribute to inflammation(Lawrence, 2009; Reece et al., 2022). This pathway plays a critical role in cell cycle regulation and is heavily involved in cancer pathogenesis due to its influence on cell proliferation and apoptosis. In the MAPK signaling pathway context, the PI3K-AKT pathway can influence cell proliferation and angiogenesis, mainly through the ERK component. The PI3K-AKT pathway’s interaction with the mTOR, JAK/STAT3, chemokine, and Toll-like receptor signaling pathways allows for a complex network of regulation and cross-talk, further expanding its role in various cellular processes. Pathogen-associated molecular patterns (PAMPs) can directly influence TLR2/4 and activate the small GTPase Rac1. This activation triggers the PI3K, producing PIP3, a crucial second messenger in the PI3K-AKT pathway. PIP3 then stimulates the kinase AKT1, which is critical for cell survival, primarily through its influence on the MDM2 gene. Furthermore, the chaperone protein Hsp90 also activates AKT1, adding another level of regulation to this pathway. This complexity contributes to the range of cellular processes the PI3K-AKT pathway influences, reinforcing its importance in understanding disease pathogenesis, particularly in cancer and inflammatory conditions(Oeckinghaus & Ghosh, 2009; Porta et al., 2014; Sadikot et al., 2005).

**Fig. 5A.**
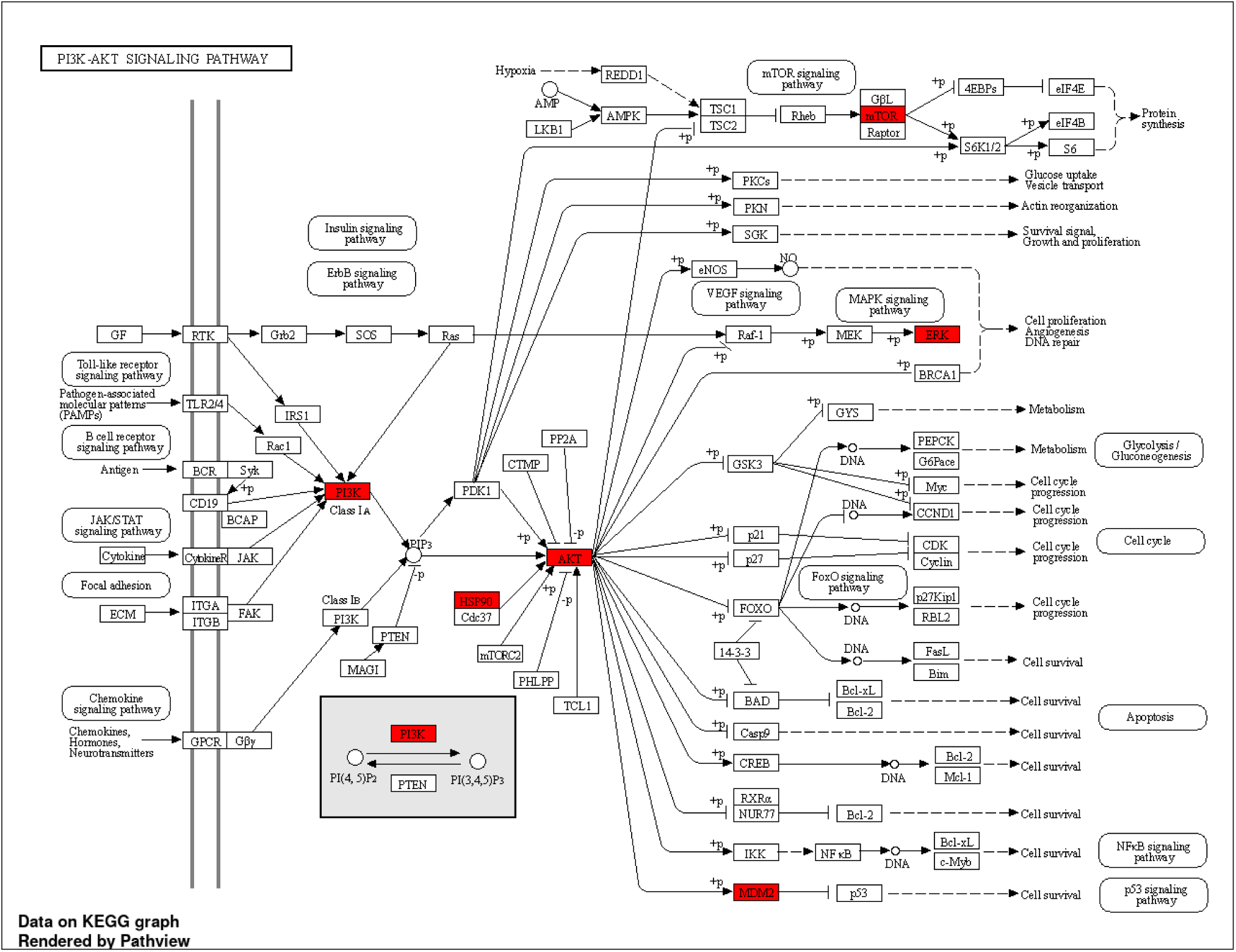
KEGG Pathway Analysis of the Top 10 Targets, with Special Emphasis on PI3K-AKT signaling pathway.

The pathway “Proteoglycans in cancer” (KEGG:05205) has a higher negative p-value of 11.70, making it the most significant path in the dataset. Proteoglycans are a group of glycosylated proteins mainly present in the extracellular matrix. They play crucial roles in many biological processes, including cell proliferation, migration, and angiogenesis, all of which are integral to cancer development and progression (**See** Fig. 5B and 6)(Ahrens et al., 2020). Several genes from data, such as AKT1, SRC, STAT3, MAPK3, and PIK3CA, are implicated in this pathway, indicating a potential role in cancer-related processes. The “Thyroid hormone signaling pathway” (KEGG:04919) is the second most significant pathway, with a negative p-value of 11.31. The thyroid hormone signaling pathway regulates metabolism, growth, and development. It involves several critical genes from the data set, including SRC, AKT1, and PIK3CA. Dysregulation in this pathway may lead to various disorders, ranging from developmental issues to metabolic diseases and certain cancers(Y. C. Liu et al., 2019). The pathway “EGFR tyrosine kinase inhibitor resistance” (KEGG:01521) also shows high significance with a negative p-value of 10.36. EGFR, a key receptor tyrosine kinase, regulates cellular activities, including proliferation and survival (**See Fig. 6**). EGFR mutations often result in over-activated EGFR pathways, causing uncontrolled cell growth, which is common in various cancers like NSCLC. EGFR tyrosine kinase inhibitors (TKIs) can hinder tumor growth by inhibiting EGFR’s tyrosine kinase activity.

**Fig. 5B.**
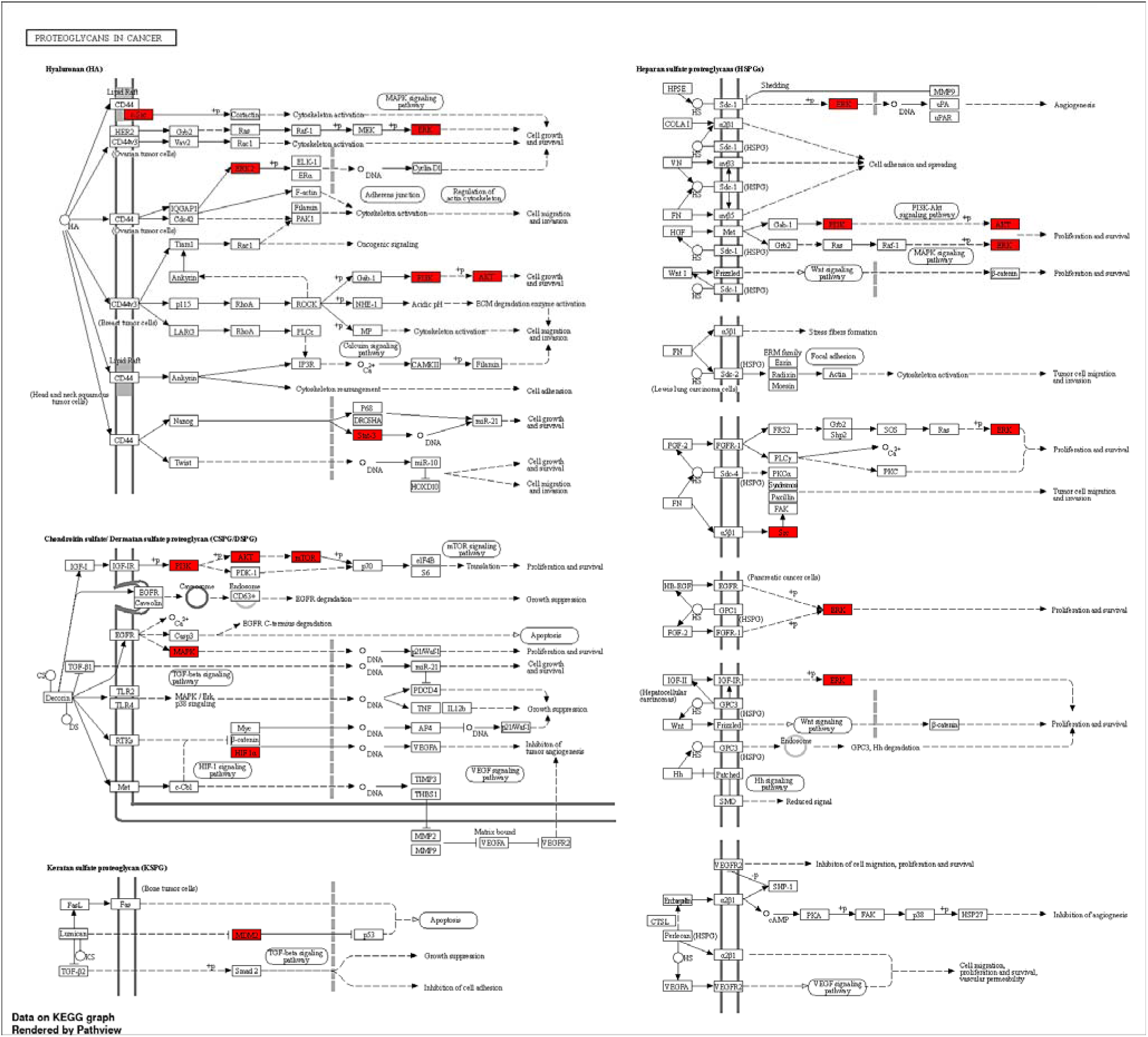
KEGG Pathway Analysis of the Top 10 Targets, with Special Emphasis on Proteoglycans in cancer.

**Fig. 6.**
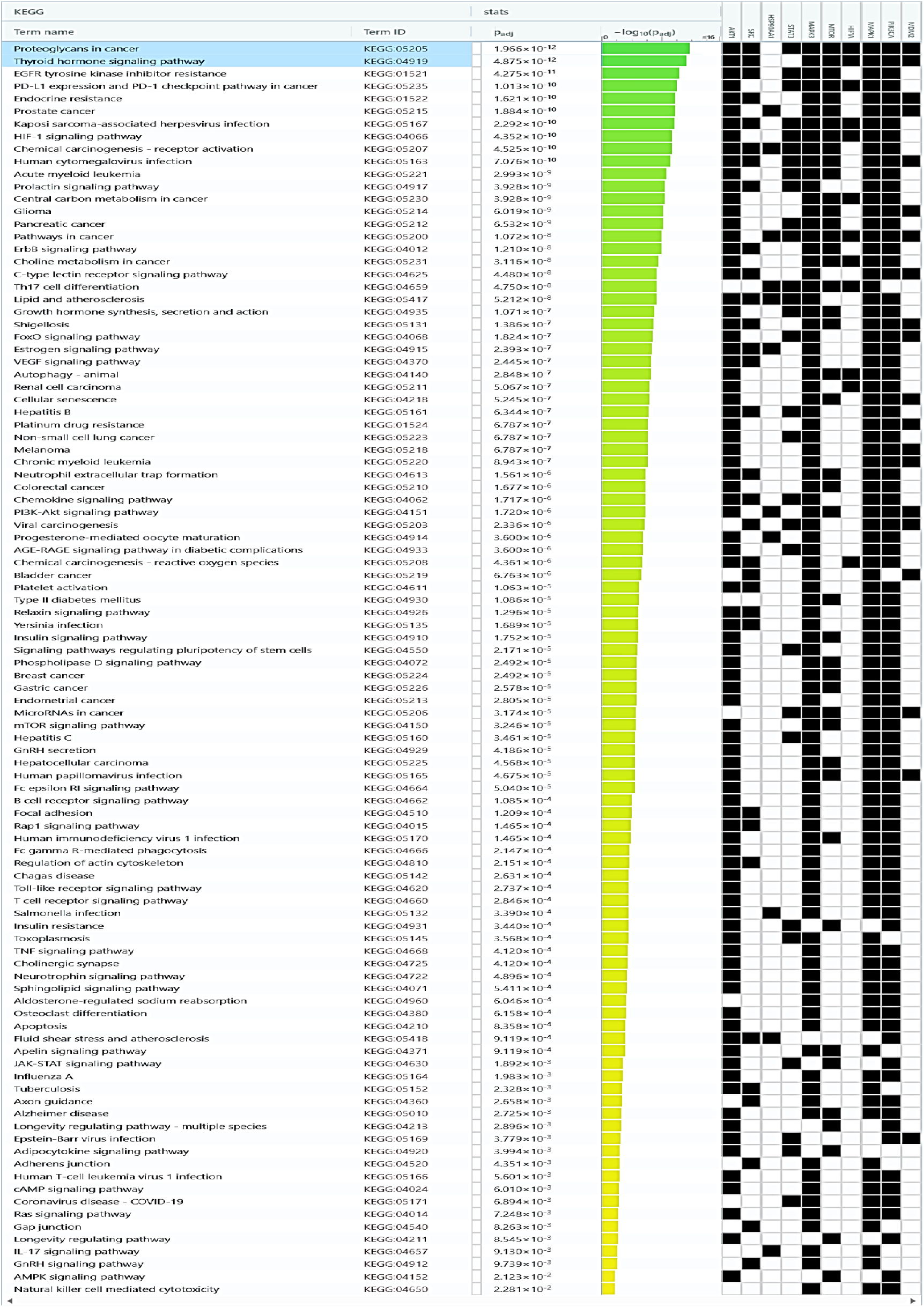
The analysis of KEGG pathways, along with their corresponding Predictive p-values and the genes they interact with.

However, resistance to these drugs often develops through mechanisms like secondary EGFR mutations or changes in other growth factor receptors. Key genes in data, such as EGFR, AKT1, PIK3CA, ERBB2, MET, and FGFR1, can contribute to EGFR TKI resistance, either by direct alterations in EGFR or by influencing related signaling pathways(Ji et al., 2013). Specific genes, like AKT1, MAPK3, MAPK1, PIK3CA, etc., appear frequently across many routes. These genes could be essential nodes in biological networks and serve as potential targets for broad-spectrum treatments.

### 3.6. Machine learning QSAR analysis

#### 3.6.1. Exploratory Data Analysis

In AKT1 gene data, After data pre-processing and curation and the elimination of missing values, 2876 compounds were selected for study. Then, all 2876 compounds were subjected to exploratory data analysis. The exploratory data analysis of a dataset of compounds revealed a greater proportion of active compounds than inactive ones. In addition, it was discovered that the pIC_50_ values of active compounds ranged from 6 to 10, whereas those of inactive compounds ranged from 3.30 to 5. The Mann−Whitney U test was performed to evaluate the statistical significance between active and inactive groups of compounds. After the U test, all of the five properties have statistical significance. Inactive molecules generally have slightly smaller MW and fewer NumHDonors than active group molecules. Active molecules tend to have larger pIC_50_, MW, and NumHAcceptors values. LogP values remain identical between active and inactive molecules (**See Fig. 7**). In SRC gene data, after data pre-processing and curation and the elimination of missing values, 3177 compounds were selected for study. Then, all 3177 compounds were subjected to exploratory data analysis. The exploratory data analysis of a dataset of compounds revealed a greater proportion of active compounds than inactive ones. In addition, it was discovered that the pIC_50_ values of active compounds ranged from 6 to 10.45, whereas those of inactive compounds ranged from 1 to 5. The Mann−Whitney U test was performed to evaluate the statistical significance between active and inactive groups of compounds. After the U test, all of the five properties have statistical significance. Inactive molecules typically possess slightly elevated LogP values compared to active group molecules. Active molecules exhibit higher pIC_50_ values than those in the inactive group. However, the values for MW, NumHDonors, and NumHAcceptors remain nearly indistinguishable between active and inactive molecules (**See Fig. 7**).

**Fig. 7.**
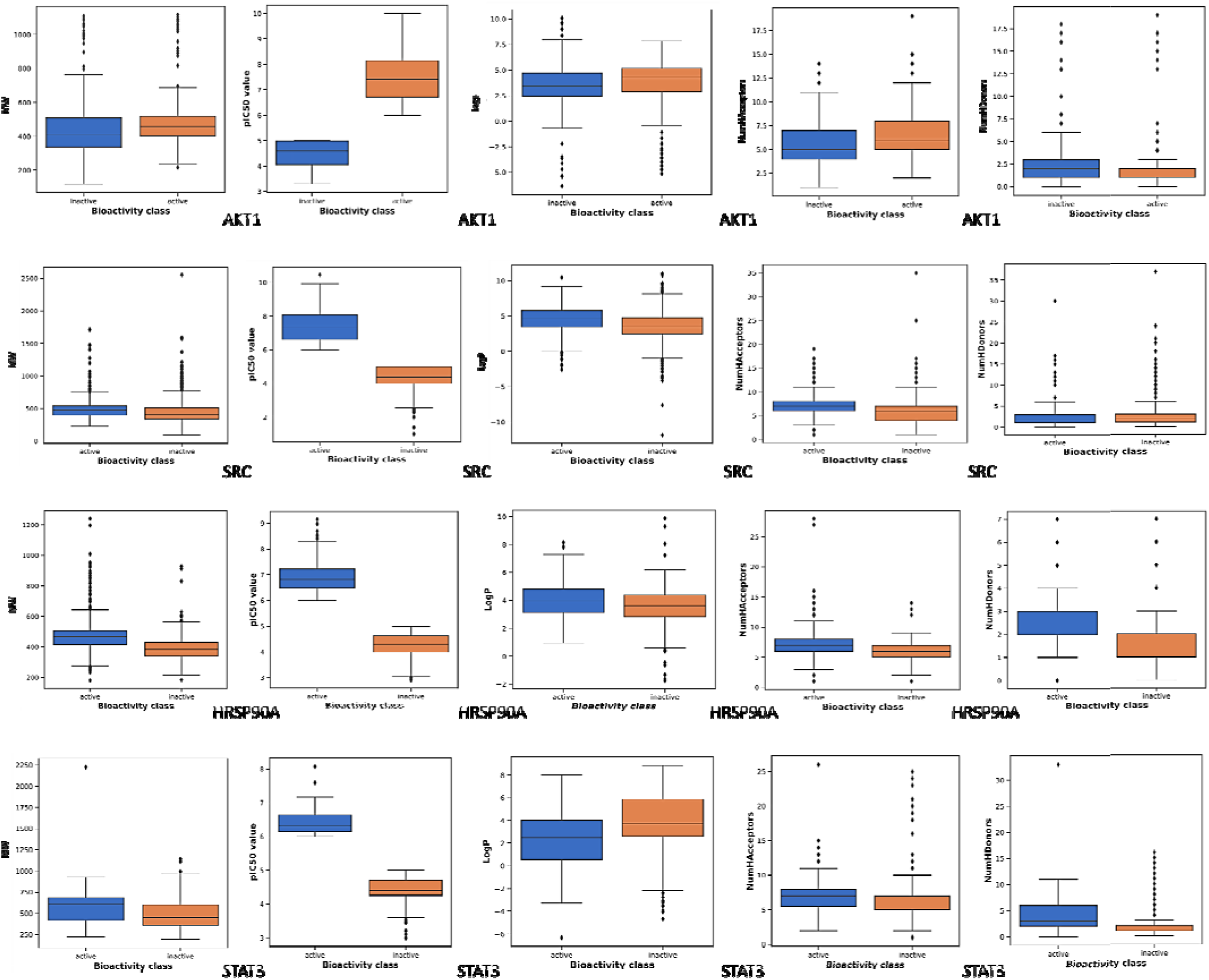
Exploratory data analysis for four genes, AKT1, SRC, HSP90A, and STAT3 inhibitors’ dataset from the ChEMBL database.

In HSP90A gene data, after data pre-processing and curation and the elimination of missing values, 1009 compounds were selected for study. Then, all 1009 compounds were subjected to exploratory data analysis. The exploratory data analysis of a dataset of compounds revealed a greater proportion of active compounds than inactive ones. In addition, it was discovered that the pIC_50_ values of active compounds ranged from 6 to 9.15, whereas those of inactive compounds ranged from 2.90 to 5. The Mann−Whitney U test was performed to evaluate the statistical significance between active and inactive groups of compounds. After the U test, all of the five properties have statistical significance. Active molecules typically show slightly elevated LogP and NumHAcceptors compared to inactive group molecules. They also tend to have higher values for MW, pIC_50_, and NumHDonors than inactive molecules (**See Fig. 7**). In STAT3 gene data, after data preprocessing and curation and eliminating missing values, 640 compounds were selected for study. Then, all 640 compounds were subjected to exploratory data analysis. The exploratory data analysis of a dataset of compounds revealed a greater proportion of active compounds than inactive ones. In addition, it was discovered that the pIC_50_ values of active compounds ranged from 6 to 8.07, whereas those of inactive compounds ranged from 3 to 5. The Mann−Whitney U test was performed to evaluate the statistical significance between active and inactive groups of compounds. After the U test, all of the five properties have statistical significance. Active molecules typically exhibit significantly higher pIC_50_ values and a greater number of NumHAcceptors than molecules in the inactive group. The values for MW, LogP, and NumHAcceptors remain nearly unchanged between active and inactive compounds (**See Fig. 7**).

#### 3.6.2. Machine learning QSAR model predication

The dataset of compounds underwent descriptor calculation using PaDELPy in the PaDEL software. Before this, PubChem fingerprints were generated individually, resulting in 881 PubChem fingerprint attributes. For developing a Random Forest-based QSAR model, Data pre-treatment GUI 1.2 was used to remove constant descriptors based on correlation coefficient and variance scores. After excluding the shared biological activity attribute, the final list of attributes for PubChem fingerprints of AKT1, SRC, HSP90A, and STAT3 were 213, 241, 253, and 262, respectively. The Kennard Stone algorithm was employed for dataset division, and the training and evaluation sets were split in an 80:20 ratio. During the training phase, QSAR models were and STAT3 (504 instances). In the subsequent testing phase, the models were evaluated using separate instances: 531 for AKT1, 576 for SRC, 186 for HSP90A, and 120 for STAT3.To generate the intended QSAR models, each descriptor dataset’s training and test datasets were loaded into the machine learning software WEKA. Information regarding the training set and testing of PubChem fingerprint and CHEMBL molecules for each target gene can be found in **Supplementary file s3**.

WeKa’s model analysis for the AKT gene shows strong predictive performance during training and testing, as indicated by the high correlation coefficient of 0.9802 and 0.8083. The model’s predictions of training are relatively close to the actual values, with a mean absolute error of 0.2225 and a root mean squared error of 0.2937. The relative absolute error of 20.1691% suggests that the model’s predictions deviate from the actual values by around 20% on average. The root relative squared error of 21.3556% indicates the variability of prediction errors. The mean absolute error and root mean squared error from test set data show 0.5868 and 0.7371, respectively. The relative absolute and root relative squared errors are around 58-59%. Cross-validation results fall between the training and testing performance, with a correlation coefficient of 0.8634. The mean absolute error and root mean squared error are 0.5299 and 0.702, respectively. The relative absolute and root relative squared errors are approximately 48% and 51%, respectively. These results suggest that the model generalizes reasonably well to new data. For the SRC gene, the QSAR model achieves a high correlation coefficient of 0.9867 during training, similar to HSP90A.

The mean absolute error and root mean squared error are 0.2227 and 0.3, respectively. The relative absolute and root relative squared errors are 14.8335% and 17.3816%, respectively. The model maintains strong performance during testing, with a correlation coefficient of 0.9147. The mean absolute error and root mean squared error are 0.5065 and 0.6198, respectively. The relative absolute and root relative squared errors are around 36%, suggesting good generalization. Cross-validation results also show high predictive ability, with a correlation coefficient of 0.8983. The mean absolute error and root mean squared error are 0.5677 and 0.7648, respectively. The relative absolute error and root relative squared error are approximately 37%, indicating consistent and reliable performance. The model analysis for the HSP90AA gene demonstrates high predictive accuracy during training, with a correlation coefficient of 0.9867. The mean absolute error and root mean squared error are 0.1403 and 0.1993, respectively. The relative absolute and root relative squared errors are 16.1236% and 17.1802%, respectively. The model maintains strong performance during testing, with a correlation coefficient of 0.9295. The mean absolute error and root mean squared error are 0.3371 and 0.4213, respectively. The relative absolute and root relative squared errors are around 37%, indicating good generalization. Cross-validation results also show high predictive ability, with a correlation coefficient of 0.9011. The mean absolute error and root mean squared error are 0.3553 and 0.5073, respectively. The relative absolute error and root relative squared error are approximately 40%, suggesting consistent performance across different folds.

For the STAT3 gene, the QSAR model achieves a high correlation coefficient of 0.9713 during training, indicating predictive solid ability. The mean absolute and root mean squared errors are 0.1719 and 0.2541, respectively, implying accurate predictions. The relative absolute and root relative squared errors are 23.079% and 27.1873%, respectively. However, on the test set, the model’s performance slightly decreases, with a correlation coefficient of 0.783. The mean absolute error and root mean squared error increase to 0.3219 and 0.4605, respectively. The relative absolute and root relative squared errors are around 59% and 70%, respectively. Cross-validation results show a correlation coefficient of 0.7102, suggesting good performance compared with training. The mean absolute error and root mean squared error are 0.4488 and 0.6589, respectively. The relative absolute and root relative squared errors are approximately 60% and 70%, respectively. The robustness of the QSAR models was inferred from the high correlation coefficients observed in both the training and test sets, suggesting a high degree of reliability. Additionally, the outcomes of tenfold crossvalidation for each model demonstrated a notable level of satisfaction, further affirming the models’ performance (**See Fig.8A**).

**Fig.8.**
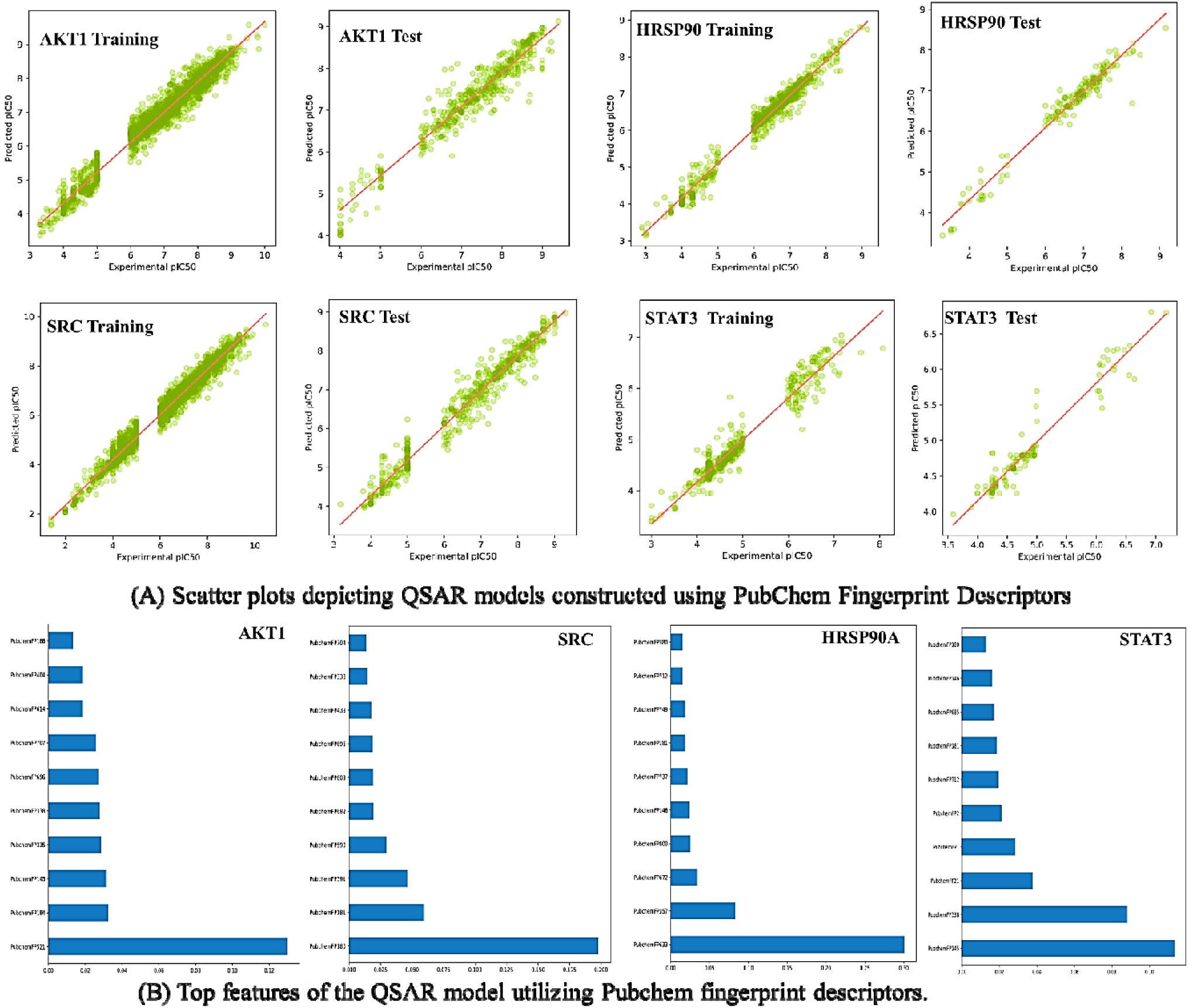
Scatter Plots of QSAR Models Utilizing Pubchem Fingerprint Descriptors for Training and Test Sets, and VIP Plot Illustrating the Key Features of the QSAR Model Incorporating Pubchem Fingerprint Descriptors against Four Genes.

To discern the pivotal molecular fingerprints and their respective contributions to bioactivity within QSAR models, a comprehensive feature importance analysis was conducted. This investigation involved the utilization of the Random Forest regressor algorithm to pinpoint the top ten molecular fingerprints for each QSAR model. The Variance Importance Plots (VIP) were generated using the matplotlib package in Python, providing a visual representation of the significance of these fingerprints (**See Fig.8B**.). The most significant descriptors in the Pubchem fingerprint-based model were identified as follows: PubchemFP521 (C:N-C-[#1]) in AKT1, PubchemFP180 (containing at least one saturated or aromatic nitrogen-containing ring of size 6) in SRC, PubchemFP633 (N-C-C:C-C) in HSP90, and PubchemFP145 (including at least one saturated or aromatic nitrogen-containing ring of size 5) and PubchemFP338 (C(∼C)(∼C)(∼H)(∼N)) in STAT3. For the PubChem fingerprints-based model targeting the AKT1 gene, the VIP analysis highlighted PubchemFPs 143, 184, 186, 335, 338, 404, 521, 614, 696, and 707 as the most influential molecular fingerprints. Similarly, in the context of the SRC gene, the VIP plot identified PubchemFPs 180, 181, 338, 391, 439, 590, 609, 682, 696, and 704 as the key contributors to bioactivity. Moving to the HSP90A gene, the VIP analysis underscored the significance of PubchemFPs 146, 181, 357, 380, 633, 672, 712, 737, 749, and 800. Lastly, within the context of the STAT3 gene, PubchemFPs 1, 2, 21, 145, 146, 180, 181, 338, 685, and 712 were identified as the critical molecular fingerprints. Based on feature selection, structural insights for the best descriptor-containing compounds were investigated for both models individually (**See Fig. 8B)**.

In the context of AKT1, specific analysis has revealed that clinical drugs 443654 (CHEMBL379300), CHEMBL3899716, and CHEMBL3966806 exhibit consistent fingerprints associated with distinct molecular features. These fingerprints include PUbchemFP143 (greater than or equal to 1, any ring size 5) and PUb-chemFP521 (C:N-C-[#1]). Experimentally determined pIC_50_ values for these compounds were 9.796, 10, and 9.824, respectively. For SRC, quantitative structure-activity relationship (QSAR) data analysis was conducted on the VIP plot. The FDA-approved drug DASATINIB (CHEMBL1421) and Chembl IDs CHEMBL1241676 and CHEMBL196797 were observed to possess common PubChem fingerprints. These fingerprints, specifically PubchemFPs 180 (greater than or equal to 1 saturated or aromatic nitrogen-containing ring size 6), 181 (greater than or equal to 1 saturated or aromatic heteroatom-containing ring size 6), and PubChem Fp696 (C-C-C-C-C-C-C-C), were reflected in experimental pIC_50_ values of 9.301, 9.921, and 9.824. These findings suggest particular structural attributes contributing to the compound’s bioactivity. In the case of HSP90A, QSAR data analysis of the VIP plot revealed shared Pubchem fingerprint attributes in FDA approved drugs REBLASTATIN (CHEMBL267792), BIIB021 (CHEMBL467399), LUMINESPIB (CHEMBL252164), and Chembl IDs CHEMBL2205798, CHEMBL4873718, and CHEMBL2205245 (**see Fig. 9**.). The common characteristics include PubChem146 (greater than or equal to 1 saturated or aromatic heteroatom-containing ring size 5), PubChem181 (greater than or equal to 1 saturated or aromatic heteroatom-containing ring size 6), PubChem357 (C(∼C)(:C)(:N)), and PubChem633 (N-C-C:C-C). The corresponding experimental pIC_50_ values are 8.30, 8.29, 8.10, 9.15, 9.14, and 9, reinforcing the structural attributes responsible for their bioactivity **(Fig.9**.).

**Fig.9.**
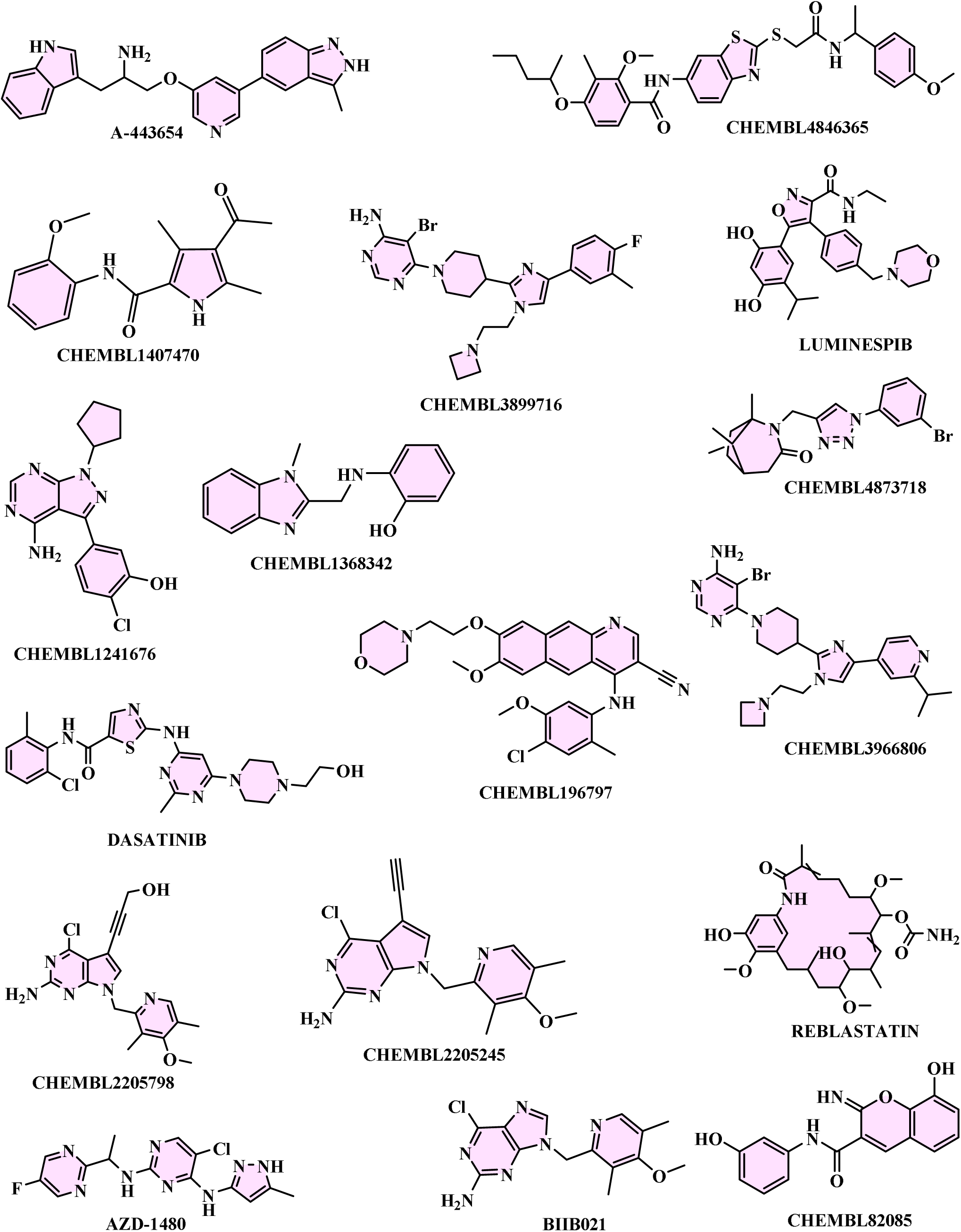
Identification of structural insights for PubChem Fingerprint Descriptors through analysis of top-performing molecules.

Lastly, in STAT3, a QSAR analysis of the VIP plot was performed for FDA-approved drug AZD-1480 (CHEMBL1231124) and Chembl IDs CHEMBL1368342, CHEMBL1407470 and CHEMBL4846365. Shared PubChem fingerprints were identified, such as PubChem146 (greater than or equal to 1 saturated or aromatic nitrogen-containing ring size 5), PubChem146 (greater than or equal to 1 saturated or aromatic heteroatom-containing ring size 5), PubChem181 (greater than or equal to 1 saturated or aromatic heteroatom-containing ring size 6), PubChem357 (C(∼C)(:C)(:N)), and PubChem633 (N-C-C:C-C). The experimental pIC_50_ values were measured at 7.097, 8.071, 7.593, and 7.17, further elucidating the structural attributes that contribute to the bioactivity of these compounds **(Fig.9**. and **Table 3**). Information regarding the training set and testing of PubChem fingerprint and CHEMBL molecules for each target gene can be found in **Supplementary file s3.**

**Table 3.** Interpretation for the most significant PubChem and substructure fingerprints.

| Best Features | Interpretation |
| --- | --- |
| PubchemFP1 | $\geq 8$ H |
| PubchemFP2 | $\geq 16$ H |
| PubchemFP21 | $\geq 8$ O |
| PubchemFP143 | $\geq 1$ any ring size 5 |
| PubchemFP145 | $\geq 1$ saturated or aromatic nitrogen-containing ring size 5 |
| PubchemFP146 | $\geq 1$ saturated or aromatic heteroatom-containing ring size 5 |
| PubchemFP180 | $\geq 1$ saturated or aromatic nitrogen-containing ring size 6 |
| PubchemFP181 | $\geq 1$ saturated or aromatic heteroatom-containing ring size 6 |
| PubchemFP184 | $\geq 1$ unsaturated non-aromatic heteroatom-containing ring size 6 |
| PubchemFP186 | $\geq 2$ saturated or aromatic carbon-only ring size 6 |
| PubchemFP335 | C(~C)(~C)(~C)(~H) |
| PubchemFP338 | C(~C)(~C)(~H)(~N) |
| PubchemFP357 | C(~C)(:C)(:N) |
| PubchemFP380 | C(~O)(~O) |
| PubchemFP391 | N(~C)(~C)(~C) |
| PubchemFP404 | N(:C)(:C)(:C) |
| PubchemFP439 | C(-C)(-N)(=O) |
| PubchemFP521 | C:N-C-[#1] |
| PubchemFP590 | C-C:C-O-[#1] |
| PubchemFP609 | Cl-C-C-N-C |
| PubchemFP614 | C-C-O-C-C |
| PubchemFP633 | N-C-C:C-C |
| PubchemFP672 | O=C-C=C-[#1] |
| PubchemFP682 | O-C-C-C-C-N |
| PubchemFP685 | O=C-C-C-C-N |
| PubchemFP696 | C-C-C-C-C-C-C-C |
| PubchemFP704 | O=C-C-C-C-C-C-C |
| PubchemFP707 | O=C-C-C-C-C(N)-C |
| PubchemFP712 | C-C(C)-C(C)-C |
| PubchemFP737 | Cc1cc(N)ccc1 |
| PubchemFP749 | Nc1cc(N)ccc1 |
| PubchemFP800 | CC1CC(N)CCC1 |

In the context of validation parameters, a comparative analysis was conducted to assess the chemical space encompassed by the training and test sets. This evaluation involved the application of the PCA bounding box method, aiming to determine the applicability domain of the molecular fingerprint datasets developed within this study. The method’s efficacy in detecting outliers within both the fingerprint models was examined. The PCA analysis was executed during the training phase, encompassing the instances for AKT1 (2255 instances), SRC (2481 instances), HSP90A (791 instances), and STAT3 (504 instances). Subsequently, in the testing phase, distinct instances were employed for model evaluation, namely 531 for AKT1, 576 for SRC, 186 for HSP90A, and 120 for STAT3, utilizing the PubChem fingerprint dataset. The outcomes of this analysis revealed that the chemical space spanned by the test set remained within the boundaries of the chemical space occupied by the training set. Consequently, it was determined that the developed fingerprint datasets exhibited applicability domains encompassing the test set. Furthermore, an examination of the PCA scores plot indicated a significant similarity in the relative chemical space occupied by compounds within both the training and test sets, as depicted in Fig. 10.

**Fig.10.**
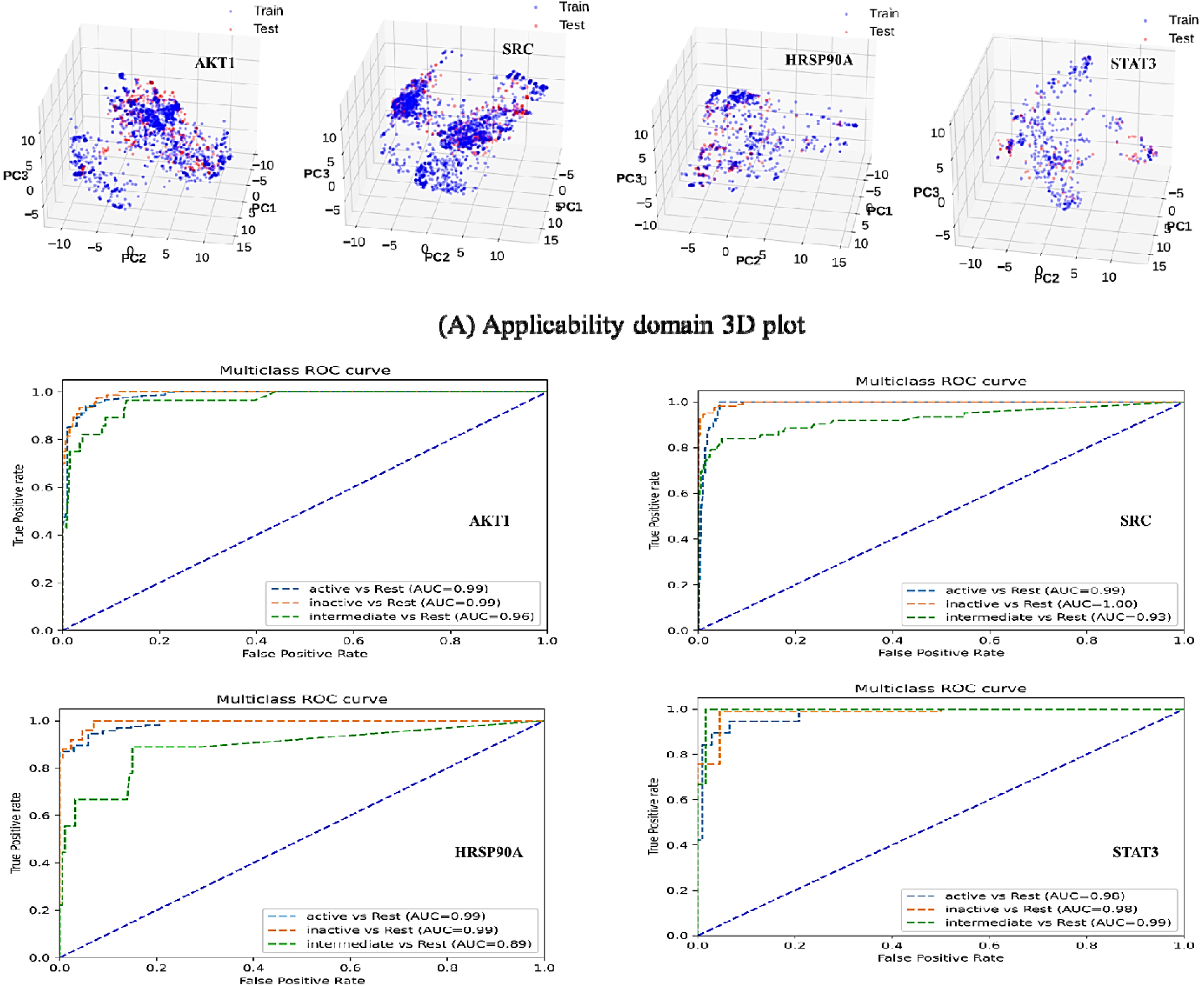
Applicability domain assessed through PCA application and ROC plot generated for PubChem fingerprint descriptor-implemented QSAR models, respectively.

The QSAR models were validated by applying Receiver Operating Characteristic (ROC) analysis, yielding pertinent insights into the predictive performance of the four target genes. Specifically, for the AKT1 gene, the computed Area Under the Curve (AUC) values were 0.99, 0.99, and 0.96 for active, inactive, and intermediate molecules, respectively. Similarly, for the SRC gene, the ROC analysis yielded AUC values of 0.99, 1.00, and 0.93 for the respective molecular classes. The HSP90A gene demonstrated AUC values of 0.99, 0.99, and 0.89 for active, inactive, and intermediate molecules. In contrast, the validation of the QSAR model for the STAT3 gene revealed AUC values of 0.98, 0.98, and 0.99 for the corresponding molecular categories. These AUC values collectively underscore the commendable and dependable performance of the QSAR models in accurately predicting molecular interactions (**See Fig.10**).

### 3.7. Prediction of bioactivity of phytochemicals using generated machine learning models

Chalcone derivatives, identified via an intensive network pharmacology screening, were assessed for bioactivity prediction using fingerprint-based machine learning models. Notably, **RA1** displayed strong interactions with Hsp90A, indicating potential as a potent inhibitor for this gene. Multi-target potential was evident in several derivatives, including **RA1, RA2,** and **RA10**, highlighting their adaptability across various pathways. Compound RA1, with its notable pIC_50_ value of 5.76 against Hsp90A, displays promising inhibitory effects, indicating its potential for diverse applications. Additionally, **RA1** exhibited substantial activity against AKT1, SRC, and STAT3 (pIC_50_: 4.89, 4.36, and 5.09), showcasing multi-target capability. Compound **RA2** exhibited significant interactions with Hsp90A (pIC_50_ = 5.62) and STAT3 (pIC_50_ = 5.09), indicating modulation potential (**See Table 4**). While interactions with AKT1 and SRC (pIC_50_ = 4.85 and 4.43) were slightly lower, **RA2’s** multi-target potential was evident. Compound **RA3** showed meaningful interactions with Hsp90A (pIC_50_ = 5.48) and STAT3 (pIC_50_ = 4.82), suggesting inhibitory effects. Interactions with AKT1 and SRC (pIC_50_ = 4.81 and 4.5) contribute to its diverse bioactivity (**See Table 4**). Compound **RA1** and **RA2** consistently exhibited higher pIC50 values, indicating relatively stronger inhibition against most target genes. In contrast, Compound **RA10** displayed lower activity across all genes. Subsequently, the chalcone derivatives underwent molecular docking studies.

**Table 4:**
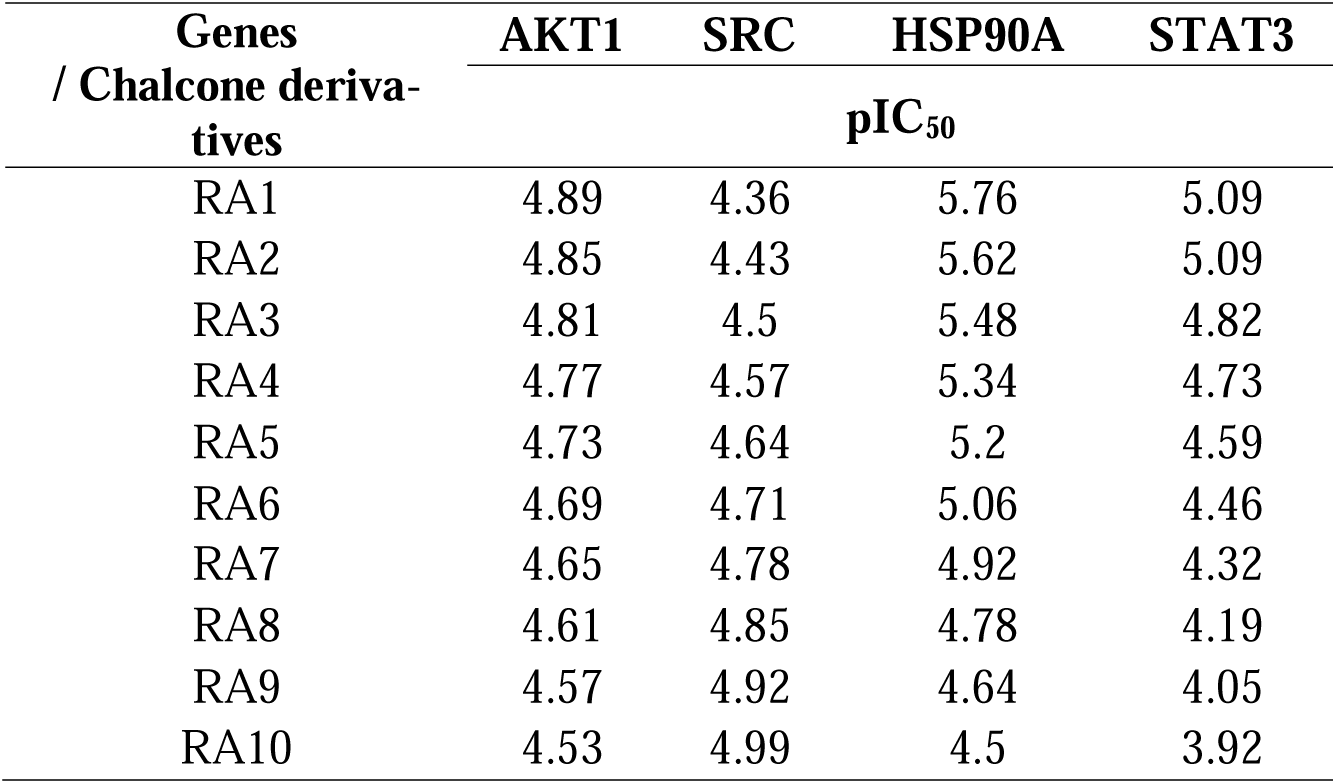
Predicted bioactivity of chalcone derivatives using generated machine learning models.

### 3.8. Web Application Development

We developed a Python-based web application named ASHS-Pred, utilizing the Streamlit library. This application leverages established molecular fingerprint-based models for AKT1, HSP90AA1, STAT3, and SRC genes. In creating the web application, various Python libraries were employed, including scikit-learn, pandas, subprocess, os, base64, and pickle. ASHS-Pred operates by considering the SMILES representations of multiple molecules and their corresponding names or IDs provided by the user within a text file. Upon uploading this text file containing molecular information, the application conducts predictions for the loaded molecules’ inhibition activity (pIC_50_) against the specified genes. The application employs established fingerprint-based random forest models to calculate the pertinent molecular fingerprints for the loaded molecules. Subsequently, the predicted activity is presented as pIC_50_ values, along with their respective molecule names.

Users can download the activity values and molecule names in CSV format directly from the application. The complete source code for ASHS-Pred is openly accessible on GitHub at the following URL: https://github.com/RatulChemoinformatics/QSAR. To utilize the application, users are required to have the Anaconda Navigator interface installed on their systems, along with Streamlit and other necessary package dependencies. The installation process is detailed in the readme file available on the GitHub repository. Following these instructions, users can accurately predict molecular activity against the four target genes using the ASHS-Pred application.

### 3.9. Molecular Docking

A molecular docking approach was employed to investigate the mechanisms underlying chalcone-based derivatives’ anti-inflammatory, antibacterial, anticancer, antidiabetic, and antifungal activities. The docking was performed against four target proteins, namely AKT1, SRC, HSP90A, and STAT3. Additionally, a set of ten chalcone derivatives and compounds with Chembl IDs were included in the study. The results of the docking analysis revealed that compounds **RA1 to RA7** exhibited superior binding affinities compared to other compounds across the four target genes. Notably, chalcone derivatives **RA1 to RA7** demonstrated comparable binding affinities to the clinical drug A-443654 (dock score = -10.9 Kcal/mol) against the AKT1 gene. Among these derivatives, Compound **RA6** displayed exceptionally high binding affinity (dock score = -10.7 Kcal/mol) towards the AKT1 target gene. Dasatinib, a known drug, exhibited significant binding affinity against the SRC target gene with a docking score of -10.5 Kcal/mol (**See Table 5**).

**Table 5.** Binding affinity scores of all the chalcone derivatives against four distinct targets.

| Compound Name | Target Genes |  |  |  |
| --- | --- | --- | --- | --- |
|  | AKT1 | SRC | HSP90A | STAT3 |
|  | Dock score (Kcal/mol) |  |  |  |
| RA1 | -10.4 | -9.8 | -10.2 | -7.4 |
| RA2 | -10.4 | -10.2 | -10.2 | -7.0 |
| **RA3** | -10.5 | -10.2 | -9.7 | **-7.5** |
| RA4 | -10.1 | -10.1 | -9.8 | -7.3 |
| **RA5** | -10.6 | **-10.7** | **-10.9** | -7.1 |
| **RA6** | **-10.7** | -10.6 | -10.5 | -7.4 |
| RA7 | -10.5 | -10.6 | -10.5 | -7.2 |
| RA8 | -8.7 | -8.3 | -8.6 | -6.1 |
| RA9 | -8.9 | -8.2 | -8.6 | -6.2 |
| RA10 | -8.7 | -8.5 | -8.6 | -6.2 |
| **A-443654** | **-10.9** | - | - | - |
| CHEMBL3899716 | -10.9 | - | - | - |
| CHEMBL3966806 | -10.8 | - | - | - |
| CHEMBL1241676 | - | -8.7 | - | - |
| CHEMBL196797 | - | -10.2 | - | - |
| CHEMBL82085 | - | -9.5 | - | - |
| **DASATINIB** | - | **-10.5** | - | - |
| BIIB021 | - | - | -8.8 | - |
| CHEMBL2205245 | - | - | -9.3 | - |
| CHEMBL2205798 | - | - | -9.1 | - |
| CHEMBL4873718 | - | - | -9.7 | - |
| **LUMINESPIB** | - | - | **-9.6** | - |
| REBLASTATIN | - | - | -7.6 | - |
| AZD-1480 | - | - | - | -6.5 |
| CHEMBL1368342 | - | - | - | -6.1 |
| CHEMBL1407470 | - | - | - | -6.1 |
| **CHEMBL4846365** | - | - | - | **-7.0** |

Interestingly, Compound **R5** showed an even better dock score of -10.7 Kcal/mol, surpassing the previously mentioned drug. Furthermore, among the studied compounds, Compound **RA5** demonstrated the strongest affinity against HSP90A with a docking score of -10.9 Kcal/mol, outperforming the FDA-approved drug Luminespib, which achieved a docking score of -9.6 Kcal/mol. Compound **R5** shows the highest docking scores for SRC and HSP90A, suggesting its potential to interact with these target genes. Compound **R6** demonstrates the highest docking score for AKT1, making it a potential candidate for targeting this gene. The docking scores suggest that Luminespib has a notable affinity for HSP90A. The docking scores for Dasatinib indicate a strong interaction with the SRC target gene. The docking scores point to a potential interaction between CHEMBL4846365 and STAT3. The docking scores for **A-443654** indicate a strong interaction with the AKT1 target gene.

In the context of AKT1, the clinical drug A-443654 does not engage in hydrogen bonding interactions. However, it establishes notable molecular interactions through pi-sigma interactions at Gln79 and Val270, alongside pi-pi stacked formations at Gln79. Additionally, alkyl and pi-alkyl interaction formations manifest at Lys268, Val270, and Trp80. In contrast, Compound **R6** forms two hydrogen bonding interactions: one with Asp292 involving the urea moiety’s NH group, and another involving an oxygen atom and the methoxy group with Arg86. A further interaction is observed with Tyr326 through van der Waals interactions. Notably, Compound **R6** exhibits alkyl and pi-alkyl interaction formations at Leu264, Leu210, and Phe55. In contrast, a pi-stacked interaction occurs at Trp80 (**See Fig.11**.). For SRC, the drug Dasatinib does not establish hydrogen bonding interactions. Nevertheless, it demonstrates molecular interactions, such as pi-sigma interactions at Met314, along with alkyl and pi-alkyl interaction formations at Val377, Val323, Ala403, Leu393, Phe405, Ala293, Val281, Ile336, and Lys295. Conversely, Compound **R5** does not display hydrogen bonding interactions but presents alkyl and pi-alkyl interactions at Val323, Ala403, Val313, His384, Val281, and pi-pi stacked formation at Phe405 (**See Fig.11**.)

**Fig.11.**
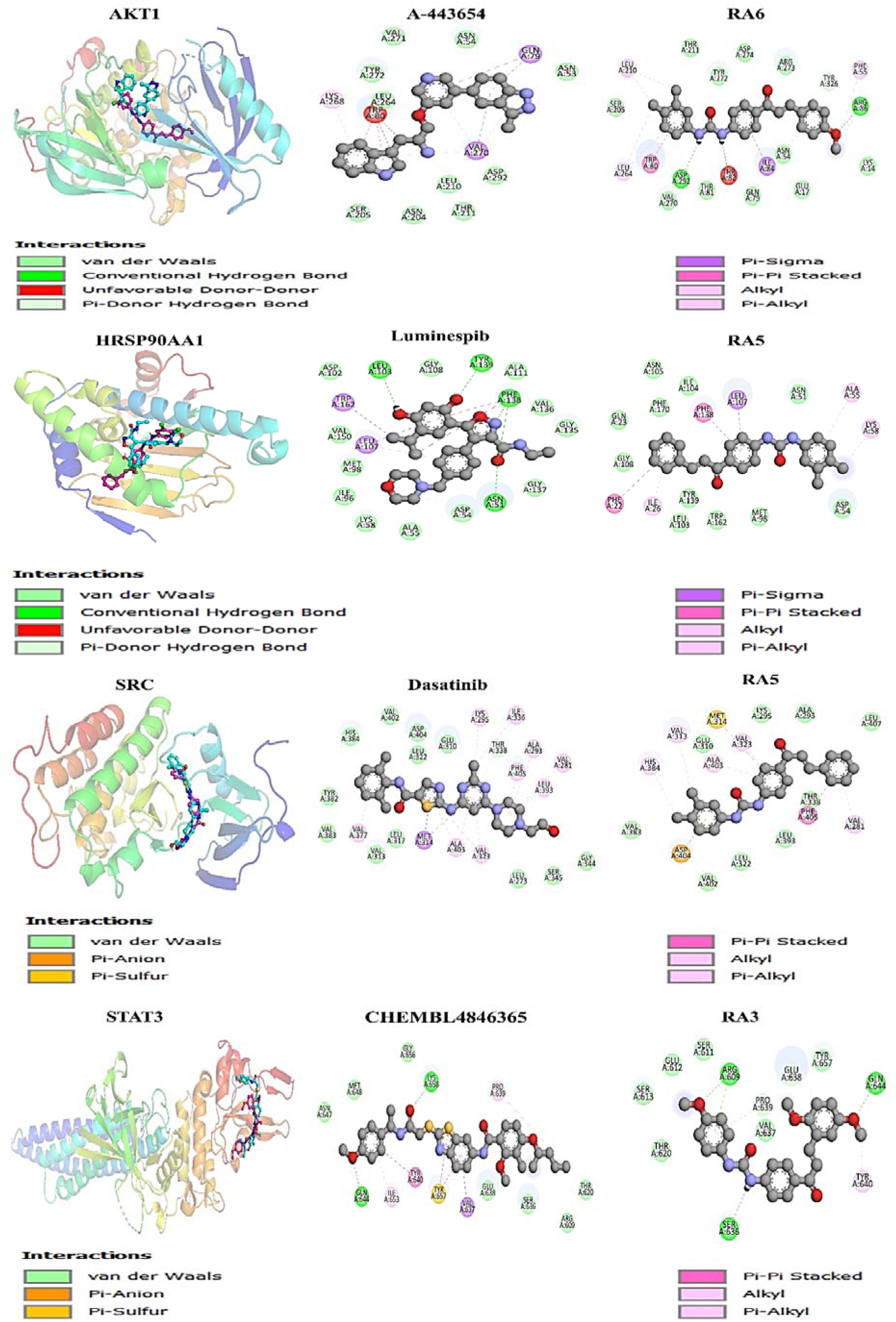
3D visualization of compound-protein interactions and 2D analysis for selected compounds (RRA5, RA6, CHEMBL4846365, Dasatinib, Luminespib, A-443654) with the protein

In the case of HSP90A, the drug Luminespib forms four hydrogen bonding interactions: the isoxazole ring’s nitrogen atom interacts with Phe138, while the carboxamide oxygen atom interacts with Asn51 and Phe138. Another interaction arises between the oxygen atom of the 4-isopropylbenzene-1,3-diol moiety and Tyr139, as well as Leu103. Further interactions include pi-sigma interactions at Trp162 and Phe138, pi-pi stacked formations at Phe138, and alkyl and pi-alkyl interaction formations at Leu107. In contrast, Compound **R5** does not engage in hydrogen bonding interactions. Still, it presents alkyl and pi-alkyl interactions at Ile26, Ala55, and Lys58, along with pi-stacked interactions at Phe22 and Phe138, and a pi-sigma interaction at Leu107 (**See Fig.11.)**. Regarding STAT3, Regarding the STAT3 target gene, Compound **RA3** exhibited remarkable binding affinity with a docking score of -7.5 Kcal/mol compared to CHEMBL4846365 (-7.0 Kcal/mol) and the clinical drug AZD-1480 (-6.5 Kcal/mol). Compound **RA3** displays the highest docking score for STAT3, indicating its potential as a candidate for targeting this gene. Compound CHEMBL4846365 forms two hydrogen bonding interactions: one with the methoxy-substituted benzene ring’s oxygen atom and Gln644 and another between the urea group’s oxygen atom and Lys658. Furthermore, pi-sigma interactions occur at Val637, while pi-pi stacked formations manifest at Tyr640 and Tyr657. Alkyl and pi-alkyl interaction formations are evident at Ile653 and Pro639. In contrast, Compound **R3** establishes three hydrogen bonding interactions: the oxygen atom of the methoxy-substituted benzene ring interacts with Arg609, while the second di-methoxy benzene-substituted ring interacts with Gln644, and the urea’s NH group interacts with Ser636. Moreover, alkyl and pi-alkyl interactions form at Tyr640 and Pro639 (**See Fig.11.)**.

### 3.10. Molecular Dynamics Analysis

A comprehensive molecular dynamics simulation running 200 nanoseconds was conducted using Desmond software to meticulously evaluate the formation of an optimal complex involving compounds RA3, RA5, RA6, CHEMBL4846365, Dasatinib, Luminespib, A-443654, and the target protein. The analysis focused on critical parameters such as root-mean-square deviation (RMSD), root-mean-square fluctuation (RMSF), and essential interactions between the protein and ligands.

The simulation results indicated that compound RA5 achieved a state of stability in terms of the RMSD values of the C-alpha atoms within the protein complex after the 10-nanosecond threshold, maintaining steady values around 2.0 Angstroms in the SRC target and 1.5 Angstroms in HSP90AA1 throughout the simulation. For the SRC target, ligand RA5 exhibited an initial equilibration phase lasting approximately 20 nanoseconds, subsequently maintaining stability within the binding pocket up to the 200-nanosecond mark (Figure 12). The RMSD of the protein also fluctuated with the ligand and, after 175 ns, slightly decreased, mirroring the initial running time from 25 ns to 172 ns. For the HSP90AA1 target, ligand RA5 displayed the same stable profile, with an upward trend towards stability between 12 and 172 nanoseconds, showing steady RMSD values around 1.5 Angstroms post the initial equilibration phase of 10 nanoseconds. After 175 ns, the ligand showed smaller fluctuations until 200 ns with RMSD values around 1.7 Angstroms. Meanwhile, the protein also showed less fluctuation throughout, remaining within 3.5 Angstroms from 25 to 172 ns, but after that, it showed a conformational shift and slightly increased until 200 ns **(Figure 12)**. Moreover, the RMSD of the known drug Luminespib initially fluctuated until 120 ns after which it stabilized and remained stable from 130 to 200 ns against the HSP90AA1 gene. Dasatinib shows that the RMSD of the protein backbone of all the complexes stabilized at approximately 1.5 Angstrom before 100 ns of simulation and then from 125 to 150 ns it increased and became more fluctuated; however, after 150 ns, Dasatinib became stable throughout the period against the SRC gene.

**Fig.12.**
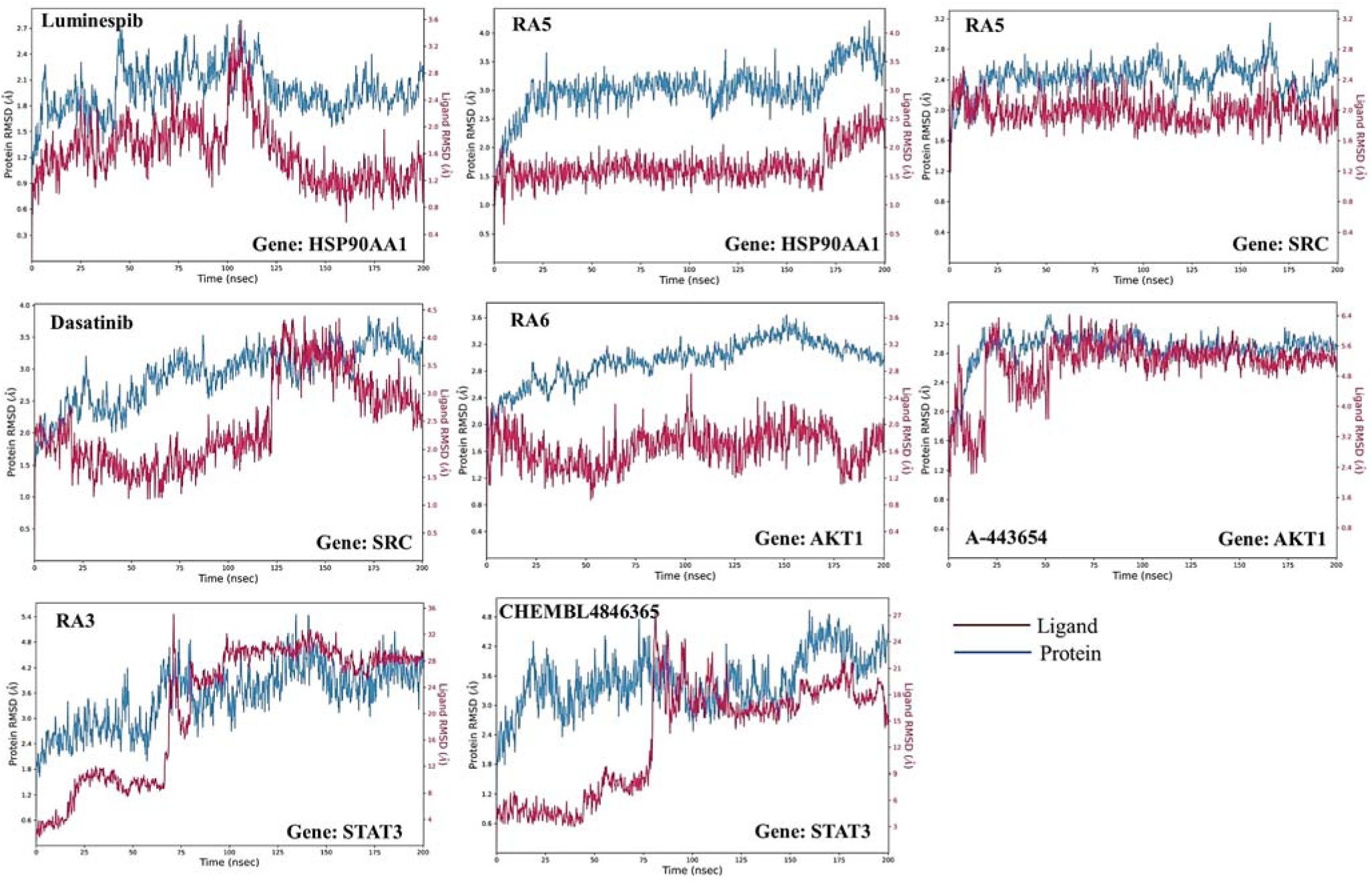
Analysis of the Root Mean Square Deviation (RMSD) of the hit compounds obtained from molecular docking studies against the target gene through Molecular Dynamics (MD) simulation.

RMSF values for compound RA5 for both targets, SRC and HSP90AA1, highlighted significant fluctuations primarily in the protein’s loop and terminal regions, while lower RMSF values at the binding site indicated stable interactions between the protein and ligands. Additionally, the secondary structural composition of the protein was analysed. For compound RA5 against the HSP90AA1 target, the structural elements, including alpha-helices and beta-strands, constituted 46.35% of the protein’s structure, thereby contributing to its structural stability and functional efficacy. Specifically, helices and strands accounted for 25.51% and 20.84% of the total structure, respectively. In the case of SRC, these elements comprised 39.76% of the protein’s structure, with helices and strands representing 26.34% and 13.42%, respectively (**Figure 13**).

**Fig.13.**
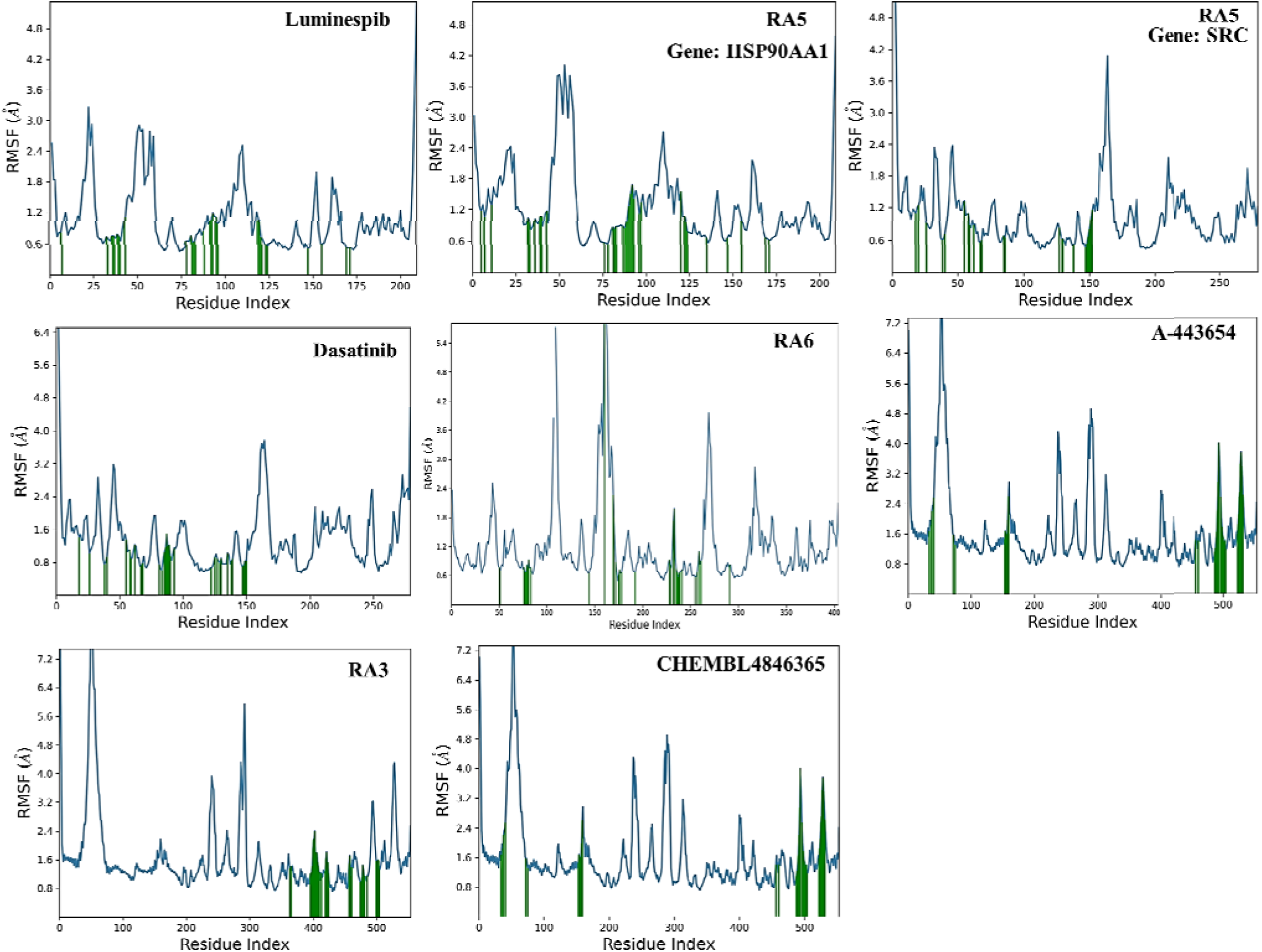
Analysis of the Root Mean Square Fluctuation (RMSF) of the hit compounds obtained from molecular docking studies against the target gene through Molecular Dynamics (MD) simulations.

The detailed analysis further explored the interactions between the ligands and the protein’s amino acid residues, illustrated through a histogram plot in **Figure 14**. This plot clearly shows the different types of interactions - hydrogen bonding (marked in green), water bridges (in blue), and hydrophobic interactions (in purple), highlighting their importance in the binding process. Compound RA5 exhibited four hydrogen bonds against HSP90AA1, particularly with amino acids Tyr139 (oxygen atom of the urea group with 91%), Leu103 (two hydrogen bonds, NH atom of the urea group with 96% and 99%), and Phe138 (with a water molecule, and those water molecules interact with the oxygen atom). A di-substituted chlorobenzene ring interacted with Phe170 residue as a hydrophobic interaction with 37%. On the other hand, Compound RA5 exhibited three hydrogen bonds against SRC, particularly with amino acids Asp404 (oxygen atom of the urea group with 92%), Glu310 (two hydrogen bonds with two water molecules, and those water molecules interact with the NH atom of the urea group with 37% and 41%). A di-substituted methoxy-containing benzene ring interacted with Phe405 residue as a hydrophobic interaction with 53% (**See Supplementary File**).

**Fig.14.**
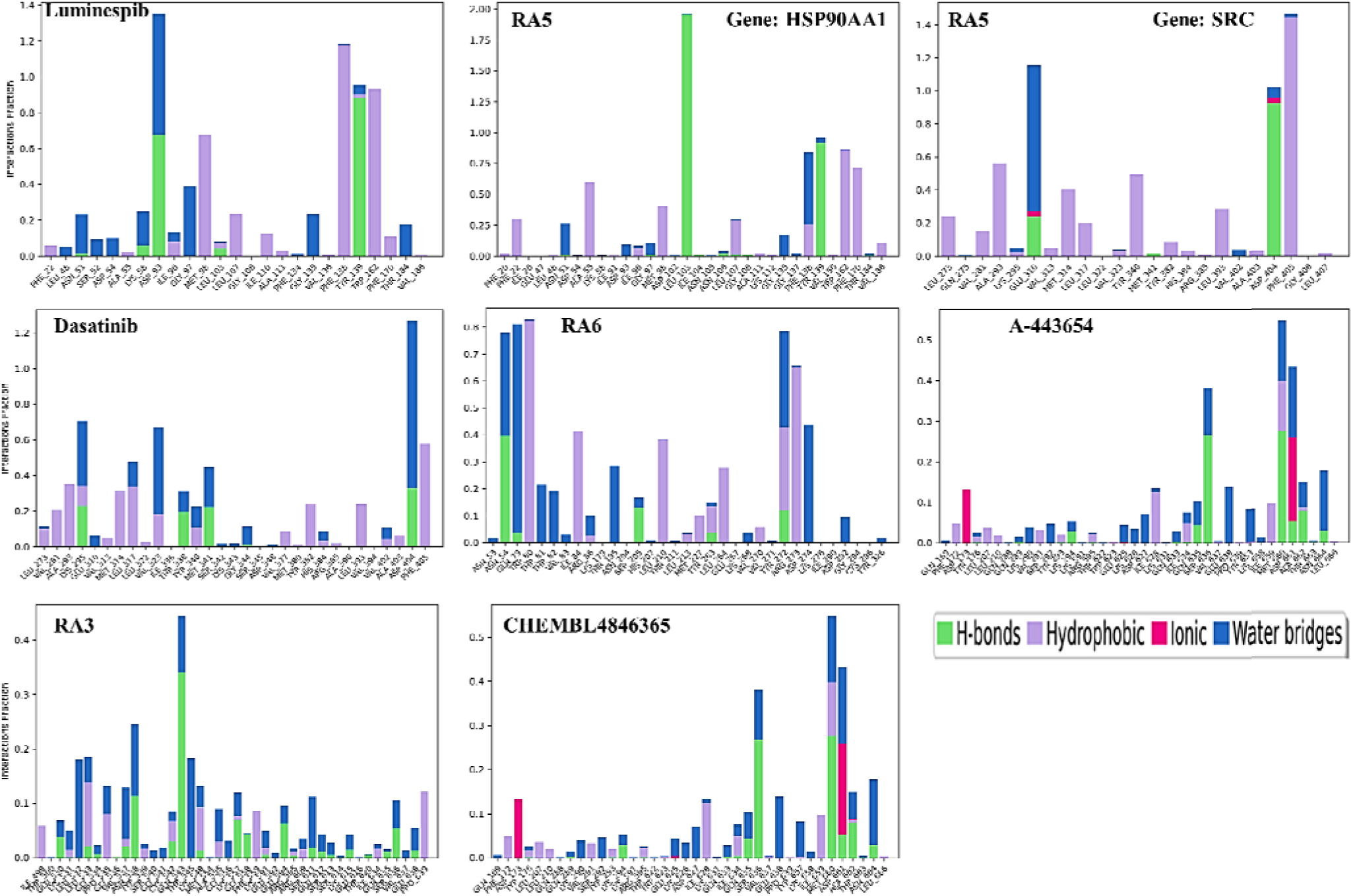
Analysis of the 2D Histogram of Protein-Ligand Contact for the Hit Compounds Derived from Molecular Docking Studies Against the Target Gene via Molecular Dynamics (MD) Simulations.

Against the AKT1 gene, the clinical drug A-443654 demonstrated stability in RMSD values, reaching 2.8 Angstroms, higher than compound R6. Compound R6 maintained stability over time within 1.6 Angstroms, while A-443654 showed more consistent stability after 70 ns up to 200 ns, exhibiting steady RMSD values (**Figure 12**). RMSF values for both compounds highlighted significant fluctuations primarily in the protein’s loop and terminal regions, while lower RMSF values at the binding site indicated stable interactions between the protein and ligands. Overall, both compound R6 and drug A-443654 displayed similar secondary structural composition in the protein, with structural elements, including alpha-helices and beta-strands, constituting 40.52% of the protein’s structure. This composition contributes to its structural stability and functional efficacy, with helices and strands accounting for 18.50% and 22.01% of the total structure, respectively. The detailed analysis further explored the interactions between the ligands and the protein’s amino acid residues, as illustrated through a histogram plot in **Figure 14**. Compound R6 exhibited all hydrogen bonding with water molecules, with 54% of interactions with Gln79 facilitated by the oxygen atom attached to the benzene ring, and 36% and 30% of interactions with Asp274 and Tyr272, respectively, facilitated by the NH atom of the urea group. Asn54 directly interacted with the oxygen atom of the urea group by 39% and connected with water molecules by 37%, which in turn are connected with the oxygen atom. Arg273 and Trp84 were linked with di-substituted chloro and methoxy-containing benzene rings through pi-cation and hydrophobic interactions, at 64% and 43%, respectively. Conversely, drug A-443654 displayed only two hydrogen bonding interactions: one from the NH group of the pyridine ring with Ser205 at 36% and the other with Asn53 from the NH group of the benzimidazole ring at 45% (**See Supplementary File)**.

Against STAT3, as illustrated in **Figure 12**, the RMSD plot for compound CHEMBL4846365 shows the protein and ligand RMSD over time. Initially, both protein and ligand RMSD values rise, typical as the system equilibrates. After this period, the ligand RMSD stabilizes, indicating that the compound has found a relatively stable conformation within the binding site. However, the protein RMSD continues to exhibit some fluctuations, suggesting that while the ligand may be stable, the protein is still undergoing conformational changes, possibly adjusting to the ligand’s presence or due to its dynamic nature. In the case of compound R3, the RMSD plot depicts a change over time, maintaining stability in the binding pocket from 100 ns to 150 ns, with some conformational shifts observed around the 10 to 100 nanosecond range, followed by stability from 125 to 150 ns (**Figure 12**). The protein’s RMSD, while fluctuating, does not show a pronounced rise, implying a more rigid structure or less conformational change in response to ligand binding compared to the complex with compound CHEMBL4846365. RMSF values, demonstrated in **Figure 13**, exhibit significant fluctuations mainly in the protein’s loop and terminal regions, with lower RMSF values at the binding site suggesting stable interactions. The structural elements of compound R3, including alphahelices and beta-strands, constituted 57.15% of the protein’s structure, contributing to its structural stability and functional efficacy, with helices and strands accounting for 40.63% and 16.51%, respectively. In contrast, for CHEMBL4846365, these elements comprised 57.43% of the protein’s structure, with helices and strands representing 40.42% and 17.02%, respectively. Compound R3 exhibited one hydrogen bonding interaction with the oxygen atom of the urea group by Gln543 at 33%. Conversely, compound CHEMBL4846365 did not show any significant contribution to interaction with the STAT3 gene target. This comprehensive interaction analysis underscores the specificity and diversity of ligand-protein interac ions and emphasizes the role of molecular dynamics simulations in uncovering intricate details of binding mechanisms, invaluable in the rational design of therapeutics for optimizing ligand efficacy and specificity.

#### Principal Component Analysis (PCA)

Principal Component Analysis (PCA) provides a detailed view of the interaction dynamics between diverse compounds and their target proteins throughout Molecular Dynamics (MD) simulations (Figure 15). This technique captures key aspects of the compounds’ stability and the range of their motion when bound to protein targets. In the case of Luminespib against the HSP90AA1 gene, the PCA plot shows data points tightly grouped near the origin for both principal components. This clustering signifies a consistent interaction dynamic, with the compound maintaining a stable conformation throughout the simulation process. Conversely, Dasatinib displays a distinct pattern when bound to the SRC protein, with data points scattered more widely along the principal component one (PC1) axis. This spread indicates a broader range of conformational states that Dasatinib may adopt during its interaction, implying a higher degree of flexibility and dynamic behavior in its binding conformations. RA5 presents an interesting case; when tested against both HSP90AA1 and SRC targets, there is a noticeable dispersal along the PC1 axis for each. This observation suggests that RA5 can induce diverse conformational states within these protein complexes. Notably, hen bound to SRC, the spread along the principal component two (PC2) axis is relatively constrained, hinting that while RA5 may exhibit a variety of shapes, these conformations likely change within a limited range within the multidimensional conformational landscape.

**Fig. 15.**
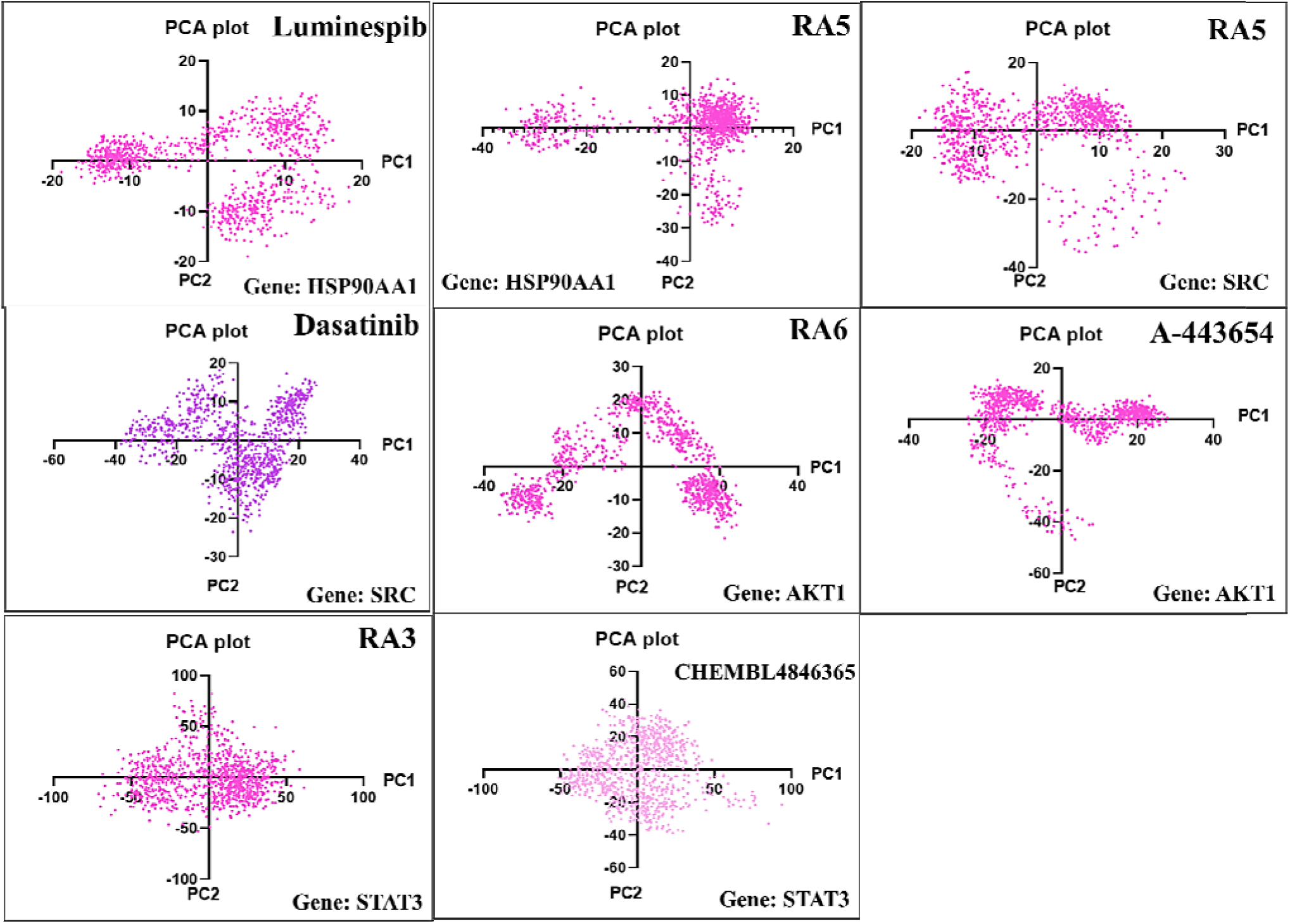
Principal Component Analysis (PCA) of Hit Compounds in Protein-Ligand Complexes.

Examining the interactions of RA6 and A-443654 with AKT1 further expands our understanding. Both compounds share a pattern of greater distribution along PC1 than PC2, which may allude to a significant conformational diversity that unfolds along a particular dimension of the interaction. Compound A-443654 shows a pronounced distribution along PC2 as well, suggesting that it can move through an even more varied range of conformations, possibly affecting different domains of the AKT1 protein. The interaction of RA3 with STAT3 is characterized by the widest distribution, especially along PC1, indicating that RA3 might access a considerable array of conformational states **(Figure 15)**. This wide range might represent various modes of binding or a high degree of structural flexibility within the ligand when it is assoc ated with the protein. CHEMBL4846365 engagement with STAT3 is also depicted with substantial spread along PC1, which is indicative of notable conformational dynamics. However, its moderate dispersal along PC2, particularly when contrasted with RA3, suggests that the diversity of its conformational changes might be less extreme across the entire structure of the complex.

#### MMGBSA

The Molecular Mechanics Generalized Born Surface Area (MM-GBSA) methods have been applied to calculate the free energy of binding for a series of compounds against various gene targets, providing insghts along with contributions from Coulombic, covalent, hydrogen bonding, lipophilic, packing, self-contact, solvation, and van der Waals interaction. For compound A443654 targeting AKT1, the MM-GBSA binding energy is notably high at -64.31 kcal/mol, with significant contributions from lipophilic interactions at - 22.74 kcal/mol and van der Waals forces at -59.68 kcal/mol. This suggests a substantial nonpolar interaction component, complemented by Coulombic interactions at -11.06 kcal/mol. Similarly, CHEMBL4846365 against STAT3 shows a binding energy of -31.80 kcal/mol, with a relatively lower van der Waals contribution of -30.78 kcal/mol, indicating a slightly less hydrophobic interaction compared to A443654 with AKT1. The lipophilic interactions for CHEMBL4846365 are also lower at -8.58 kcal/mol. Dasatinib’s binding to SRC is characterized by a binding energy of -83.46 kcal/mol, with a large negative contribution from lipophilic interactions at -29.06 kcal/mol and a significant van der Waals term at -73.20 kcal/mol, reflecting strong hydrophobic and van der Waals interactions within the binding site. Luminespib shows a strong affinity for HSP90A1 with a binding energy of -85.86 kcal/mol. The notable lipophilic and van der Waals contributions of -26.50 and -62.68 kcal/mol, respectively, highlight the compound’s strong hydrophobic binding character.

For RA3 against STAT3, the binding energy is -45.08 kcal/mol. This is paired with a hydrogen bond contribution of -0.57 kcal/mol and a notable van der Waals term of -36.51 kcal/mol, indicating a good balance of polar and nonpolar interactions. Compound RA5 targeting HSP90A1 exhibits a particularly strong binding energy of -96.26 kcal/mol, with the highest lipophilic contribution among the compounds at -34.71 kcal/mol and a substantial van der Waals component at -66.31 kcal/mol, suggesting a potent interaction with the protein. RA6 interacting with AKT1 has a binding energy of -66.30 kcal/mol. Its lipophilic and van der Waals contributions are significant at -28.07 and -66.11 kcal/mol, respectively, indicative of favorable hydrophobic interactions. Lastly, RA5 against SRC shows the most potent binding energy of -100.01 kcal/mol within this dataset. The lipophilic term is extremely high at -38.98 kcal/mol, coupled with a large van der Waals contribution of -75.37 kcal/mol, which could be reflective of a tight and efficient binding to the active site.

## 4. Discussion

The focus of our recent study was to identify key chalcone compounds and Chembl libraries aimed at AKT1, SRC, HSP90A, and STAT3. These targets, by potentially inhibiting their metabolic pathways, were chosen for their capacity to act against cancer, diabetes, fungal infections, inflammation, and bacterial infections. This versatility makes chalcones a valuable candidate for drug development because they can target multiple disease pathways. The approach to achieving this objective involved the integration of machine learning, molecular mechanisms, and systems biology techniques. This approach involved structure-based high-throughput screening of small molecule databases targeting these four genes, followed by pharmacokinetic screening and docking. We also identified potential small molecule inhibitors that could block the binding sites of these target gene pathways using machine learning-assisted Quantitative Structure-Activity Relationship (QSAR) modeling and web-based affinity prediction. A multifaceted approach such as this demonstrates the potential of integrated computational methodologies for advancing the discovery and development of drugs. Thorough exploratory data analysis (EDA) in the initial data analysis played a pivotal role in shaping the subsequent phases of the research. We meticulously curated and preprocessed the dataset, unveiling significant distinctions between active and inactive compounds across various molecular properties. The Mann-Whitney U test underscored the statistical significance of these differences, underscoring the potential of chalcone derivatives as bioactive compounds.

Central to our investigation was the creation of Random Forest-based QSAR models for each target gene. These models exhibited commendable predictive performance, characterized by high correlation coefficients and acceptable error rates during training and testing. Significantly, the feature importance analysis uncovered specific molecular descriptors crucial for predicting bioactivity, offering vital insights into the structural determinants of chalcone derivatives’ effectiveness. Extensive analysis uncovered a promising group of compounds, particularly **RA1 to RA7**, which demonstrated exceptional bioactivity against the target genes. For instance, compound **RA1** displayed a remarkable pIC_50_ value of 5.76 against HSP90A, positioning it as a standout candidate for further exploration. This correlation between compound structure and bioactivity emphasizes the potential utility of chalcone derivatives in drug discovery. Molecular docking studies further elucidated the binding interactions between chalcone derivatives and the target genes. Compounds **RA5, RA6,** and **RA7** exhibited significant binding affinities, equaling or exceeding those of existing drugs, indicating their promise as potent compounds. The detailed analysis of these binding interactions reveals the specific structural features responsible for bioactivity, aiding in a rational approach to drug design. For the benefit of the scientific community, the fingerprint-based predictive models for the top genes AKT1, SRC, HRSP90AA, and STAT3 were further deployed as the ASHS-Pred web-based application (https://ashspred.streamlit.app/) and the source codes (https://github.com/RatulChemoinformatics/QSAR) along with the data sets were made available on GitHub to encourage further extension or modification of the web server. It is important to observe that as new experimental data on the individual gene inhibitors become available, the predictive model proposed here could be continuously updated to increase its coverage and accuracy. In molecular dynamic study, particularly highlighting compound RA5’s stability with SRC and HSP90AA1 targets. This stability, evidenced by consistent RMSD values, suggests potential therapeutic efficacy. Comparatively, the differences in RMSD and RMSF values between compounds, including A-443654 and R6 against the AKT1 gene, underscore the unique interaction dynamics each compound exhibits with its target. Furthermore, detailed analyses of secondary structures and ligand-protein interactions, such as hydrogen bonding and hydrophobic contacts, offer a view of binding affinities and specificities.

Discussing the results of MMGBSA, the particularly high van der Waals and lipophilic interaction energies observed for most compounds suggest that these compounds may have substantial hydrophobic contacts within the binding sites of their respective targets, which is often a hallmark of drug-like molecules. For instance, the compound RA5 against the SRC gene exhibited the most potent binding energy at -100.01 kcal/mol, marked by the highest lipophilic contribution at -38.98 kcal/mol among the dataset, underscoring its strong affinity and specificity towards the target. This suggests that RA5 could robustly occupy the hydrophobic pockets within SRC, maximizing van der Waals contacts and potentially leading to high inhibitory activity. Moreover, the compounds targeting HSP90A1, specifically Luminespib and RA5, demonstrated high binding energies and substantial lipophilic contributions, indicating effective hydrophobic interactions that could stabilize the inhibitor within the chaperone’s binding domain. These interactions, coupled with the observed hydrogen bonds, are essential for a stable drug-protein complex, enhancing the efficacy of the drug. In contrast, the lower binding energies seen with compounds like CHEMBL4846365 against STAT3 suggest weaker interactions, which could be due to less optimal alignment within the binding pocket or insufficient hydrophobic contact, potentially leading to reduced inhibitory activity. Comparing the PCA plots collectively, it is evident that the conformational stability and flexibility of these compounds when interacting with their respective targets vary. Compounds such as Luminespib exhibit a more constrained range of motion, indicative of a stable interaction, while others like RA3 and CHEMBL4846365 demonstrate significant conformational diversity, which might correlate with multiple binding modes or interactions with the protein targets. These observations are critical for understanding the dynamic nature of protein-ligand interactions and can have implications in the optimization of these hit compounds for potential therapeutic applications.

This study concisely demonstrates the significant potential of chalcone derivatives in targeting key genes, with a focus on high-efficacy compounds RA1 to RA7. It underscores the relevance of structural factors in drug design and advocates for further experimental validation. Integrating machine learning and knowledge-base neural network insights with molecular docking simulations, the research offers a promising direction for developing treatments in anti-inflammation, antibacterial, anticancer, antidiabetic, antifungal and antituberculosis areas, potentially addressing critical medical needs and advancing drug discovery.

## 5. Conclusion

We identified significant chalcone derivatives and ChEMBL libraries targeted at AKT1, SRC, HSP90A, and STAT3. The ability of chalcones to target multiple disease pathways underscores their potential in drug development. An integrated approach, combining machine learning, molecular mechanisms, and knowledge-based neural network techniques, has advanced drug discovery. Notably, chalcone derivatives RA1 to RA7 exhibited substantial bioactivity against key target genes, with RA1 showing the most promising pIC_50_ value, particularly against Hsp90A. Docking scores corroborated these findings, with RA1 displaying robust binding affinities across all genes. Remarkably, compounds RA5, RA6, and RA7 exhibited docking scores comparable to RA1, indicating similar potential. However, a decline in activity was observed from RA8 to RA10, consistent with pIC_50_ trends. Further reinforcing these findings, comprehensive molecular dynamics simulations provided deeper insights into the dynamic interactions and stability of these compounds, particularly RA5, with target proteins SRC and HSP90AA1. The simulations of 200 nanoseconds highlighted the compounds’ stability and interaction dynamics, crucial for understanding their therapeutic potential high-lighted the compounds’ stability and interaction dynamics, crucial for understanding their therapeutic potential. The consistent RMSD values of compound RA5 after the initial equilibration phase illustrate a stable interaction with the proteins, potentially contributing to its efficacy. This dynamic analysis enhances the insights provided by static docking scores and bioactivity findings, giving a fuller understanding of how the compounds interact with their target proteins. Compared to established drugs and ChEMBL compounds, chalcone derivatives demonstrated promising results, with some outperforming known drugs in binding affinity. Specifically, compound RA5 exhibited exceptional binding affinity against HSP90A, surpassing Luminespib, an FDA-approved drug. Compound RA3 exhibited significant binding to STAT3, highlighting the potential of chalcone derivatives in a range of medical applications, evidenced by their encouraging binding scores with crucial genes. Additional research into these derivatives, encompassing both in vitro and in vivo studies, is necessary to confirm their effectiveness in treating diseases related to AKT1, HSP90AA, SRC, and STAT3. Insights from machine learning models provide a robust foundation for future research in chalcone-based small molecule binding and drug discovery.

## Declaration

### Authors’ contribution statement

Writing original draft preparation: A.M. and R.B; Writing-review & editing: A.M., R.B., S.A, S.S. A.A., S.P. and M.I.; Conceptualization: A.M., R.B., and S.A; Investigation: A.M., R.B., S.A., and A.A.; Software: A.M., and R.B.; Data curation and Formal Analysis: A.M., and R.B.; Methodology: A.M., and R.B.; Resources: A.M., R.B., S.A., and S.K.; Visualization: A.M., R.B., S.A., S.S., S.P. A.A. M.I., S.K., and B.M.; Supervision: S.A. and A.A.; Project Administration: S.A. and A.A.; Validation: A.M., R.B., S.A., S.S., S.P., A.A., S.K., B.M., and M.I.; Funding acquisition: A.A. The authors have given their approval for the final version of the manuscript before its submission for publication.

## Acknowledgement

The authors extend their sincere appreciation to the following organizations for the invaluable support and funding: 1) Finnish Cultural Foundation, 2) Tampere Tuberculosis Foundation, (Ashok Aspatwar); 3) Academy of Finland, and 4) Jane and Aatos Erkko Foundation (Seppo Parkkila).

