## Supplemental File for "Elucidating Molecular Mechanism and Chemical Space of Chalcones through Knowledge Graph and Machine Learning: A Combined Bio-Cheminformatic Pipeline"

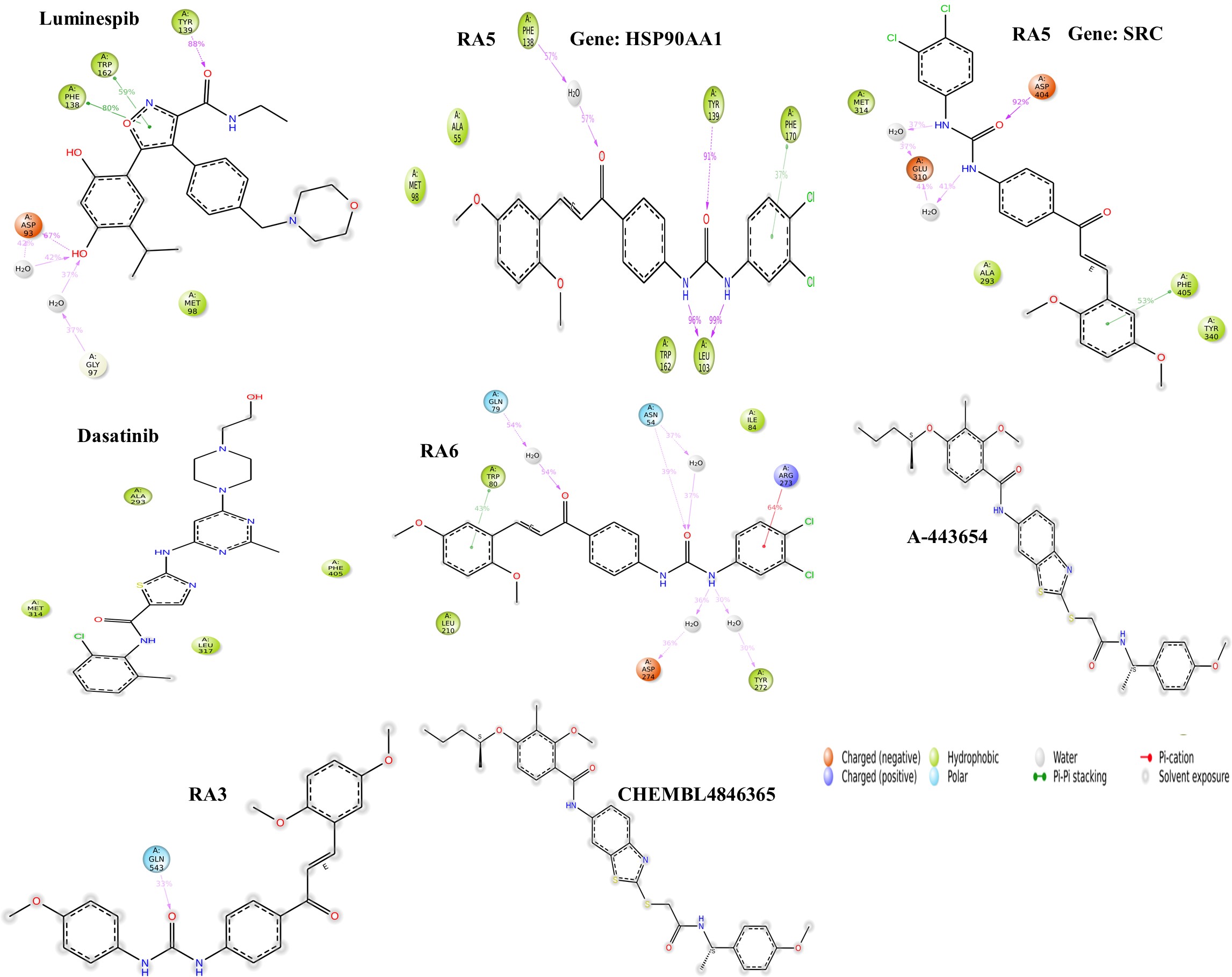

**Figure: 2D** interaction Diagram of protein and ligand through molecular dynamics study.
